# Long-term high-yield skeletal muscle stem cell expansion through staged perturbation of cytokine signaling in a soft hydrogel culture platform

**DOI:** 10.1101/2020.06.04.134056

**Authors:** Alexander M. Loiben, Kun Ho Kim, Sharon Y. Soueid-Baumgarten, Victor M. Aguilar, Jonathan Chin Cheong, Ruth F. Kopyto, Paula Fraczek, Ern Hwei Hannah Fong, Rahul Mangal, Lynden A. Archer, Benjamin D. Cosgrove

## Abstract

Muscle stem cells (MuSCs) are an essential stem cell population for skeletal muscle homeostasis and regeneration throughout adulthood. MuSCs are an ideal candidate for cell therapies for chronic and acute muscle injuries and diseases given their inherent ability to self-renew and generate progenitor cells capable of myogenic commitment and fusion. Given their rarity and propensity to lose stem-cell potential in prolonged culture, methods for *ex vivo* MuSC expansion that achieve clinical-scale stem cell yields represent a critical unmet need in muscle cell-therapeutic development. Here, we tested a microenvironment engineering approach to achieve long-term adult mouse MuSC expansion suitable for clinical demands through the combined optimization of techniques previously reported to achieve small-yield MuSC expansion in short-term cultures. We developed an optimized protocol for high-yield MuSC expansion through the combination of inflammatory cytokine and growth factor co-stimulation, temporally-staged inhibition of the p38α/β mitogen activated protein kinase signaling pathway, and modulation of substrate rigidity in long-term hydrogel cultures. We found that, on soft, muscle-mimicking (12 kPa) hydrogel substrates, a mixture of the cytokines TNF-α, IL-1α, IL-13, and IFN-γ and the growth factor FGF2 stimulated robust exponential proliferation of adult MuSCs from both wildtype and *mdx* dystrophic mice for up to five weeks of culture that was accompanied by a phenotype shift towards committed myocytes. After observing that the temporal variation in myogenic commitment coincided with an oscillatory activation of p38α/β signaling, we tested a late-stage p38α/β inhibition strategy and found that blocking p38α/β signaling after three weeks, but not earlier, substantially enhanced cell yield, stem-cell phenotypes, and, critically, preserved transplantation potential for up to five weeks of FGF2/cytokine mix culture on soft hydrogels. Notably, this retention of transplant engraftment potency was not observed on traditional plastic substrates. We estimate that this protocol achieves >10^8^-fold yield in Pax7^+^ stem cells from each starting MuSC, which represents a substantial improvement in stem-cell yield from long-term cultures compared to established methods.

**Highlights:**

- TNF-α/IL-1α/IL-13/IFN-γ cytokine cocktail supports prolonged MuSC proliferation *ex vivo* but induces differentiation.
- Cytokine cocktail regulates cell signaling with varied prolonged activation signatures.
- Effects of p38α/β inhibition on cytokine-induced MuSC expansion are stage-dependent.
- Soft hydrogels with late-stage p38α/β inhibition expand functional Pax7^+^ MuSCs long-term.

**Short summary:** Cosgrove and colleagues develop a long-term muscle stem cell expansion protocol by combining a tunable stiffness hydrogel substrate, an inflammatory cytokine cocktail, and targeted inhibition of p38 MAPK signaling. They show that soft, muscle-mimicking hydrogels with delayed p38 inhibition yield robust quantities of Pax7^+^ functional muscle stem cells.

## Introduction

Muscle stem cells (MuSCs; also called satellite cells) reside in skeletal muscle tissue in a myofiber-associated niche location and are essential for muscle homeostasis and regeneration throughout adulthood (Wang and Rudnicki, 2012). After muscle damage, quiescent MuSCs are activated and divide through self-renewal, yielding progeny that both retain a Pax7-expressing stem-cell phenotype and also differentiate into myogenic progenitor cells, which further commit into fusion-competent myocytes to repair damaged or lost myofibers (Almada and Wagers, 2016). This process of endogenous skeletal muscle regeneration involves the transient expansion of the MuSC population through self-renewal, as coordinated by a network of supporting cell types and factors, over the course of weeks, resolving in a return to stem-cell homeostatic quiescence (Bentzinger et al., 2013; Wosczyna and Rando, 2018).

Given their essential role in muscle regeneration, endogenous MuSCs and their culture progeny, a population of myogenic progenitors known as myoblasts, have been tested as a candidate for cell-based therapy for chronic muscle disease and wasting disorders, with limited clinical success (Almada and Wagers, 2016; Judson and Rossi, 2020). Notably, early-stage clinical trials using myoblasts demonstrated limited success in improving long-term muscle function to Duchenne muscular dystrophy patients, even though they resulted in detectable restoration of donor cell-derived dystrophin protein expression (Gussoni et al., 1992, 1997). These results are explained by myoblasts’ poor survival, migration, and self-renewal/expansion capacity following intramuscular transplantation, unlike primary MuSCs which are endowed with these hallmark functional capacities when isolated from healthy adult donor tissue (Bouchentouf et al., 2007; Montarras et al., 2005; Sacco et al., 2008). Though strategies have been developed to enhance transplantation outcomes for myoblasts (Borselli et al., 2011; Rao et al., 2017; Sleep et al., 2017) and myogenic stem cells derived from induced pluripotent stem cells (Rinaldi and Perlingeiro, 2014) and mesangioblasts (Price et al., 2007), muscle stem cells remain an advantageous cell therapeutic candidate due to their restricted myogenic potential and robust self-renewal capacity.

*Ex vivo* expansion of functional muscle stem cells is a bottleneck to the use of endogenous MuSCs autologous or allogeneic cell therapies for systemic muscle wasting disorders or volumetric muscle loss (Judson and Rossi, 2020). Though protocols for FACS-based cell isolation have been refined, the scarcity of MuSCs *in vivo* (only ~2-5% of all skeletal muscle cells) relative to the number of MuSCs needed for transplantation therapies necessitates a robust expansion for clinical applications. Approximately 10^3^ viable MuSCs can be isolated from a 100 mm^3^ biopsy (Blau and Webster, 1981). Though this number is sufficient to achieve functional recovery of small muscles in a dystrophic mouse model (Arpke et al., 2013), human transplantation will require a much larger number of cells. Based on prior clinical trials with myoblasts (Skuk, 2004), it is estimated that a cell therapy for DMD may require ~10^8^ functional MuSCs for individual muscles or ~10^11^ cells for whole-body therapies; this implies the need to expand *ex vivo* MuSCs isolated from muscle biopsies ~10^5^ to 10^8^-fold for therapeutic use in humans.

Given the propensity of isolated MuSCs from human and mouse tissue to differentiate and lose their stem-cell functions in traditional *ex vivo* culture platforms (Cosgrove et al., 2009; Lutolf et al., 2009), recent efforts have focused on developing biomolecular, pharmacological, and/or biophysical culture parameters to better support MuSC self-renewal and expansion outside the body. Recently, various culturing approaches have been utilized to stimulate *ex vivo* expansion of MuSCs in short- or intermediate-term cultures. Many small molecule and protein ligand factors have been shown to promote the expansion of adult mouse MuSCs, generally increasing functional Pax7-expressing cells ~2-15-fold in short-term (~1 wk) cultures (Almada and Wagers, 2016; Judson and Rossi, 2020). These approaches typically stimulate pathways promoting self-renewal or inhibit pathways driving myogenic differentiation. Reported expansion factors include the non-canonical Wnt7a ligand (Le Grand et al., 2009), the cyclic AMP activator forskolin (Xu et al., 2013), and pharmacological inhibitors of p38α/β mitogen-activated protein kinase (MAPK) (Bernet et al., 2014; Charville et al., 2015; Cosgrove et al., 2014), translational elongation factor eIF2-α (Lean et al., 2019; Zismanov et al., 2016), and the methyltransferase Setd7 (Judson et al., 2018). Furthermore, culture substrate engineering approaches have demonstrated that substrate biophysical parameters and extracellular matrix proteins can influence MuSC self-renewal *ex vivo*. Soft hydrogel platforms (typically ~10-12 kPa Young’s modulus) that mimic the rigidity of skeletal muscle tissues (which range from ~5-40 kPa, as reviewed in (Blau et al., 2015; Morrissey et al., 2016)) have shown promise for permitting MuSC self-renewal (Cosgrove et al., 2014; Gilbert et al., 2010; Lutolf et al., 2009).

Notably, Fu et al. have demonstrated that MuSCs cultured in conditioned media from activated CD3^+^ T-cells exhibit prolonged but infrequent MuSC expansion (Fu et al., 2015). Through profiling experiments, they observed that the inflammatory cytokines TNF-α, IL-1α, IL-13, and IFN-γ derived from activated T-cells were sufficient to induce MuSC expansion (Fu et al., 2015). TNF-α and IL-1α are secreted by mast cells, T-cells, and neutrophils and stimulate NFκB and p38α/β MAPK signaling (Chen et al., 2007; Egerman and Glass, 2014; Li et al., 2009; Yang and Hu, 2018). IL-13 is secreted by mast cells, T-cells, and eosinophils and predominantly activates STAT6 through type 2 innate signaling (Heredia et al., 2013; McCormick and Heller, 2015). IFN-γ is produced by T-cells, B-cells, and NK-cells and induces STAT1 and STAT3 signaling (Castro et al., 2018; Qing and Stark, 2004; Schroder et al., 2004). These pathways have been associated with both promoting and antagonizing MuSC self-renewal function, so the mechanisms by which this cytokine mixture regulates MuSC expansion remain unresolved and could be further optimized. Likewise, the standard myogenic cell mitogen FGF2, which is secreted by myofibers and myofibroblasts *in vivo*, and regulates p38α/β, MEK–ERK, AKT, and JNK signaling, exhibits a balance of self-renewal effects (Pawlikowski et al., 2017; Yablonka-Reuveni et al., 1999).

Here we tested whether long-term MuSC expansion protocols could be improved through a combined optimization of substrate biophysical parameters, expansion-promoting inflammatory cytokine factors, and targeted perturbation of related cell signaling pathways. We posited that combining these approaches may enhance the yield of functional MuSCs towards the scale needed for cell therapy applications. We show that, on soft, muscle-mimicking (12 kPa) hydrogel substrates, a mixture of the cytokines TNF-α, IL-1α, IL-13, and IFN-γ and the growth factor FGF2 stimulate exponential proliferation of adult mouse MuSCs over one month of culture, but these cells shift to committed myocyte phenotype. We found that inhibiting p38α/β signaling, which exhibits a phased activation in these long-term cultures, after three weeks, but not earlier, substantially enhances MuSC expansion and maintains stem cell-like transplantation potential. This new protocol integrating soft synthetic hydrogels with cytokine-stimulation and staged pathway perturbation achieves a ~10^8^-fold expansion in Pax7^+^ MuSCs over one month of culture, a significant improvement from established methods.

## Results

### Physiologically relevant substrate stiffness differences influence negligible effects on MuSC phenotype in short-term cultures

Given prior reports highlighting the importance of substrate stiffness in permitting MuSC self-renewal *ex vivo* (Gilbert et al., 2010), we hypothesized that subtle changes in substrate stiffness may influence the maintenance and expansion mouse MuSCs over a short-term (~1 wk) culture duration. We utilized a poly(ethylene glycol) (PEG) hydrogel system as a culture platform (Cosgrove et al., 2014; Gilbert et al., 2010; Lutolf and Hubbell, 2003). By varying the PEG weight percentage from 2.5% to 4.5%, we observed that the elastic stiffness (quantified by Young’s modulus through shear rheometry) of the PEG hydrogels ranged between 4.7 and 35 kPa (**Fig. S1A**). This range of moduli facilitated examination of the role of physiological stiffness on MuSC phenotype, as it covers stiffnesses inclusive of young adult (2-mo) mouse (~10-18 kPa), aged adult (18-mo) mouse (~20-30 kPa), and dystrophic adult mouse muscles (~40 kPa), as well as cultured myocytes and myotubes (Blau et al., 2015; Collinsworth et al., 2002; Cosgrove et al., 2009; Engler et al., 2004, 2006; Gao et al., 2008; Gilbert et al., 2010; Rosant et al., 2007; Stedman et al., 1991; Yin et al., 2013). We functionalized the PEG hydrogel for cell adhesion by conjugating the MuSC niche protein laminin (Sanes, 2003) via surface Michael addition reaction. We observed that the concentration of surface-conjugated laminin, as measured by immuno-chemiluminescence detection, was not affected by the PEG weight percentage (**Fig. S1B**).

Using an established FACS protocol (Sacco et al., 2008), we isolated CD34^+^ α7-Integrin^+^ MuSCs from the hindlimb muscles of adult (3-4 mo) C57BL/6J mice and cultured them in standard myogenic “growth medium” containing the myogenic mitogen FGF2 (referred to hereon as FGF) on laminin-conjugated hydrogels ranging from 5 to 60 kPa (**Fig. 1A**). After 7 days of culture, we observed no significant differences in MuSC morphology between substrates of different stiffness (**Fig. 1B**), or in proliferation index by counting cells (**Fig. 1C**). To characterize the MuSC phenotype, we analyzed expression of hallmark myogenic genes at 7 d by RT-qPCR (**Fig. 1D-G**). In the range of 5-60 kPa, we observed no statistically significant differences in expression of both *Pax7*, an essential transcription factor in quiescent and self-renewing MuSCs, and *Myod1*, a myogenic differentiation factor, between any modulus values (**Fig. 1D, F**). We observed subtle but statistically significant changes between very soft (5 kPa) and stiff (30-60 kPa) substrates in the expression of *Myf5*, a MuSC activation factor, and *MyoG*, a myogenic commitment factor (**Fig. 1D-G**). These results suggest differences in substrate stiffness are insufficient to induce substantial changes in MuSC phenotypes in short-term PEG hydrogel cultures, and that a range of muscle-mimetic (5-60 kPa Young’s modulus) substrates may provide similarly permissive environments for *ex vivo* MuSC maintenance.

**Figure 1.**
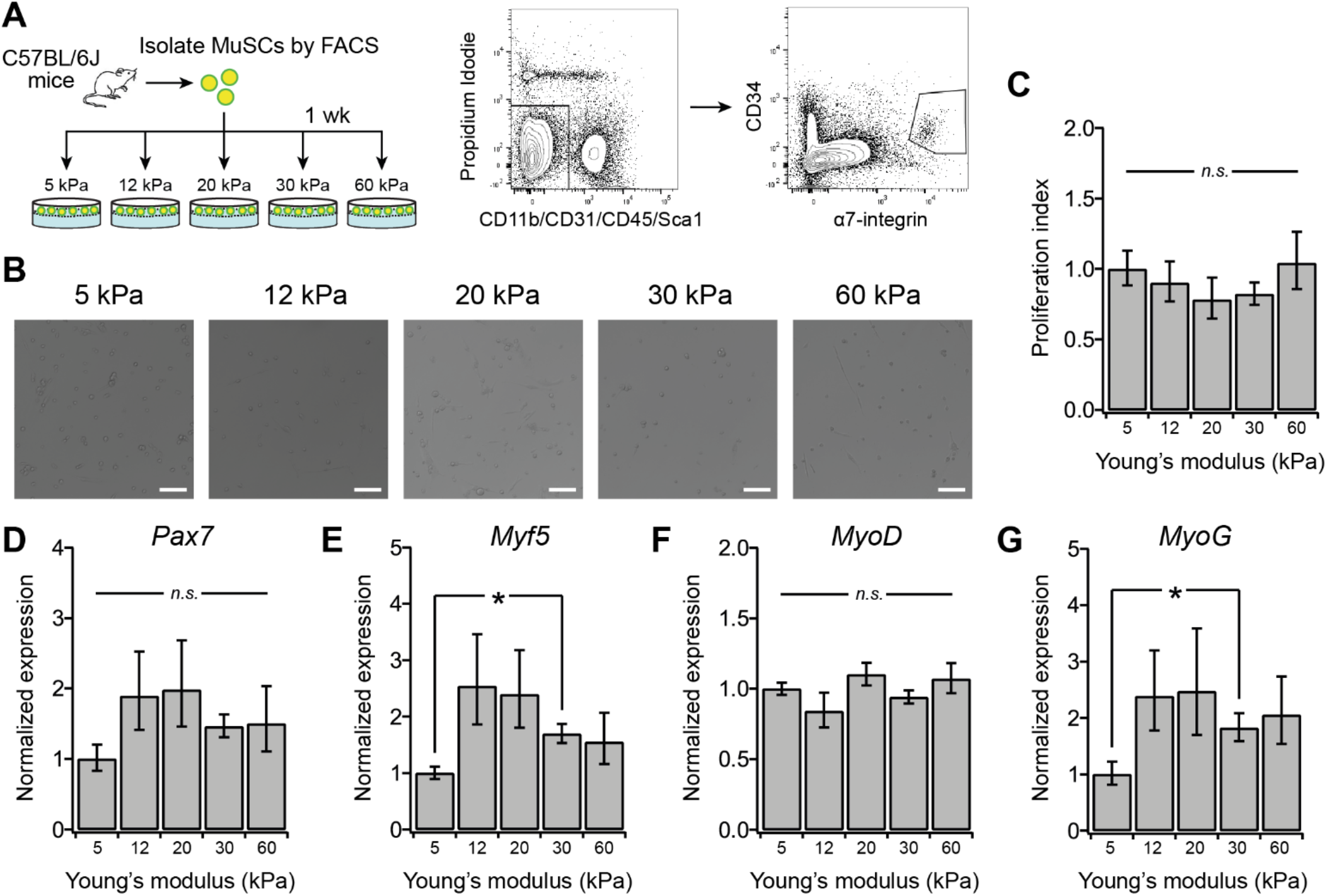
Muscle mimicking substrate stiffness differences influence negligible effects on MuSC phenotype in short-term cultures. (**A**) Experimental design. MuSC were isolated via FACS sorting for CD34^+^/α7-integrin^+^, seeded on laminin-conjugated hydrogels ranging from 5-60 kPa Young’s modulus, and treated with myogenic growth medium containing FGF2 for 1 wk. See also **Fig. S1** for hydrogel characterization. (**B**) Representative Hoffman modulation contrast images at 1 wk. Scale bar, 100 μm. (**C**) Proliferation index (cell number normalized to 5 kPa condition) after 1 wk. Mean ± s.e.m., n = 4. (**D-G**) RT-qPCR measurement of *Pax7*, *Myf5*, *Myod1*, and *Myog* expression normalized to *36B4* at 1 wk. Mean ± s.e.m., n = 4. * denotes *P* < 0.05 by Student’s T-test and n.s. denotes not significant in (**C-G**).

### Long-term cytokine treatment enhances MuSC proliferative yield while reducing stemness gene expression

Several approaches, including the inhibition of p38α/β MAPK (Bernet et al., 2014; Charville et al., 2015; Cosgrove et al., 2014) and exogenous stimulation with a combination of inflammatory cytokines TNF-α, IL-1α, IL-13, and IFN-γ (Fu et al., 2015), have been shown to promote MuSC proliferation in short- and intermediate-term cultures, respectively. We thus hypothesized that these strategies, when applied to cells maintained on soft hydrogels for long-term cultures, might synergistically enhance MuSC proliferation while maintaining their stem-cell phenotype, and thus yield an expansion of stem cell numbers. Here, we use these culture term definitions: short (1 wk), intermediate (2-3 wk), and long (4-6 wk). Given the minor differences in short-term maintenance of MuSC phenotype at 5-60 kPa (**Fig. 1**), we chose to perform long-term culture experiments on 12-kPa hydrogels as representative of soft, muscle-mimetic substrates.

We cultured adult MuSCs on laminin-conjugated 12 kPa hydrogels in growth medium containing FGF. We further supplemented the growth medium with SB203580 (a p38α/β MAPK inhibitor, hereafter referred to as p38i) and/or a mix of TNF-α, IL-1α, IL-13, and IFN-γ (hereafter referred to as “cytokine mix” or “cyt. mix”). Cells were passaged every 6 d, reseeded on new hydrogels, and tracked by extrapolating total cell counts from each passage. We detected significant differences between the conditions in total cell yield at 4 wk (**Fig. 2A**). p38α/β inhibition alone increased proliferation relative to the FGF-only controls and resulted in 10^4^-fold total cell yield relative to initial seeding. The cytokine mix had a larger proliferative effect, resulting in 10^9^-fold total yield. At 2 wk, we observed increased elongation and fusion of cells in the presence of the cytokine mix, whereas p38α/β inhibition alone did not alter cell morphology compared to the FGF-only controls, suggesting that the cytokine stimulation may promote myogenic cell activation and commitment (**Fig. 2B**).

**Figure 2.**
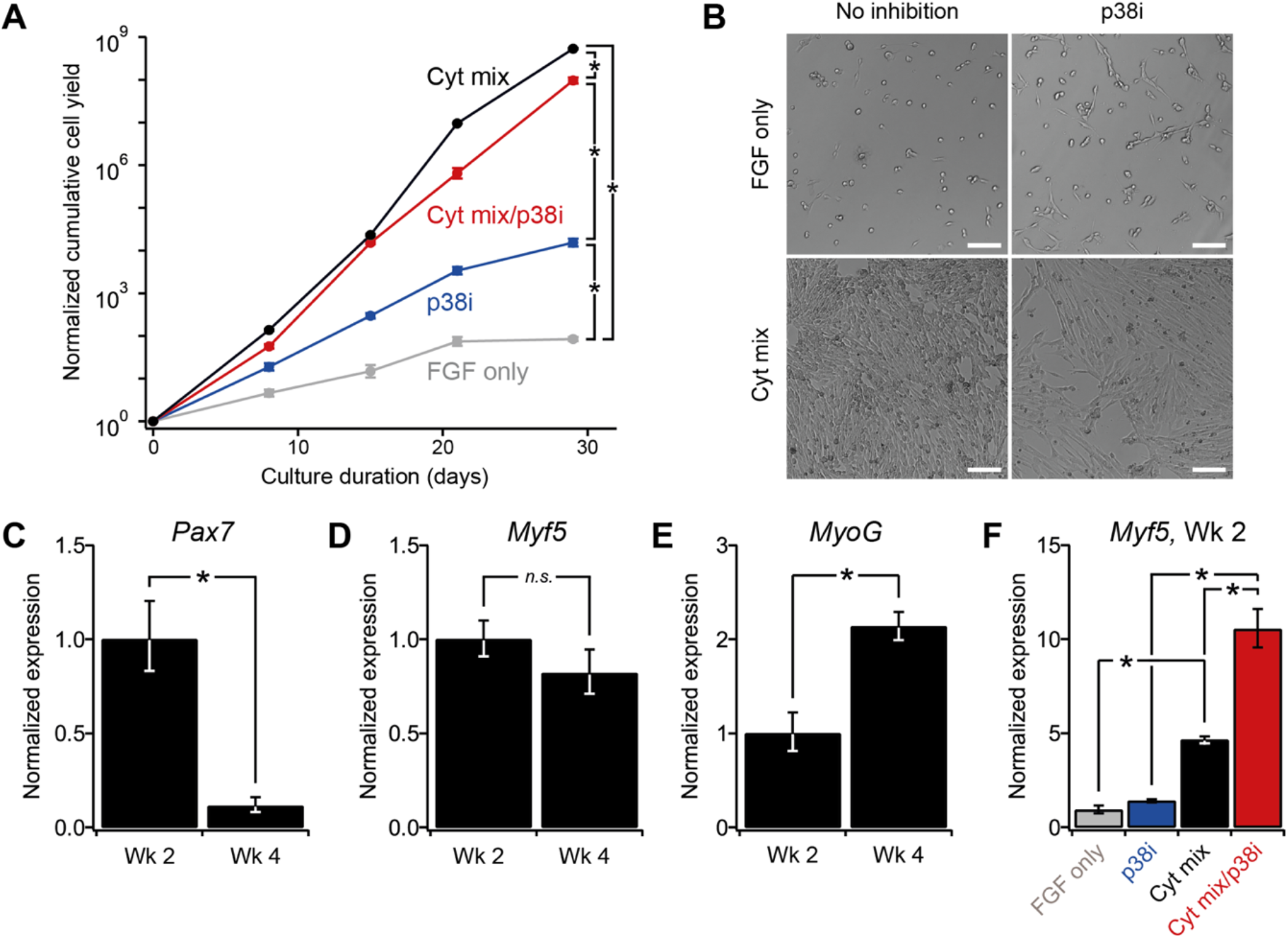
Long-term cytokine treatment enhances MuSC proliferative yield while reducing expression of stem cell-related gene. (**A-F**) Long-term MuSC cultures on 12-kPa hydrogels passaged every 6-8 d and treated with FGF2 or cytokine mix (FGF2, TNF-α, IL-1α, IL-13, and IFN-γ) and/or p38i (SB203580, 5 μM) for 4 wks. **(A**) Representative modulation contrast images at 2 wks. Scale bar, 100 μm. (**B**) Cumulative cell yield counts normalized to seeded number. Mean ± s.e.m., n = 4. * denotes *P* < 0.05 by Student’s T-test for log-transformed values at 4 wks. See also **Fig. S2A-B**. (**C**-**E**) RT-qPCR measurement of *Pax7*, *Myf5*, and *Myog* expression normalized to *36B4* in cytokine mix condition at 2 and 4 wks. Mean ± s.e.m., n = 3. (**F**) RT-qPCR measurement of *Myf5* expression normalized to *36B4* at 2 wks. Mean ± s.e.m., n = 3. * denotes *P* < 0.05 by Student’s T-test in (**C-F**).

We performed similar experiments with stiff 60-kPa hydrogels, and observed minimal differences in total cell yield at 4 wk for each of the FGF-only, p38α/β inhibition, and cytokine mix conditions between 12 kPa and 60 kPa hydrogels, indicating substrate rigidity has a negligible effect on long-term MuSC proliferation for these conditions (**Fig. S2A**). Notably, when we combined the cytokine mix with p38α/β inhibition throughout the culture duration to test if they exhibit additive effects, the total cell yield at 4 wk was 10-fold less than the yield from the cytokine mix alone (**Fig. 2B**). As such, we conclude the cytokine mix enhances proliferation throughout the long-term culture, but the effect does not synergize with long-term p38α/β inhibition. We also investigated the long-term effects of p38i and the cytokine mix on MuSCs isolated from dystrophic (*mdx*) mice. We detected increases in total cell yield similar to the wild-type control MuSCs in the presence of the cytokine mix at 4 wk, but in contrast p38i had no proliferative benefit for *mdx* MuSCs relative to FGF-only growth medium (**Fig. S2B**).

To assay the myogenic phenotype during long-term cytokine mix-stimulated hydrogel cultures, we analyzed the expression of *Pax7*, *Myf5*, and *Myog* using RT-qPCR (**Fig. 2C-F**). We found significantly lower expression of *Pax7*, negligible differences in *Myf5*, and higher expression of *Myog* at 4 wk relative to 2 wk (**Fig. 2C-E**). After 2 wk, MuSCs treated with cytokine mix expressed higher levels of *Myf5* relative to control (**Fig. 2F**) and this effect was amplified with inhibition of p38α/β. These results indicate that in long-term culture on soft hydrogels, the TNF-α/IL-1α/IL-13/IFN-γ cytokine mix induces a progressive shift towards commitment and maturation, and thus enhances proliferation at the expense of a stem-cell phenotype. The addition of prolonged p38α/β inhibition attenuates proliferative yield but preserves intermediate-term expression of *Myf5*, suggesting an enhanced activated progenitor phenotype.

### Cytokine stimulation induces time-varying activation of intracellular signaling pathways in long-term soft hydrogel culture

As summarized in **Fig. S3A**, TNF-α, IL-1α, IL-13, IFN-γ, and FGF2 stimulate the activation of varied intracellular signaling pathways in muscle stem and progenitor cells with conflicting effects on proliferation and self-renewal (Chen et al., 2007; Heredia et al., 2013; Loiben et al., 2017; Palacios et al., 2010; Qing and Stark, 2004; Yang and Hu, 2018). We hypothesized that selectively blocking individual anti-proliferative and/or pro-differentiation pathways might enhance cytokine mix-induced cell yields. To this end, we examined short-term MuSC proliferation induced by cytokine mix treatment on 12 kPa hydrogels in combination with a panel of small molecule inhibitors of the signaling mediators MEK, JNK, STAT3, p38α/β, AKT, JAK2 and IKK (**Fig. S2C**). None of these inhibitors enhanced MuSC proliferation relative to the cytokine mix by itself, and most diminished MuSC proliferation (**Fig. S2C**), suggesting multiple downstream pathways (JAK2–STAT3, p38α/β, AKT, and IKK–NFκB) are all critical for the net pro-proliferative effect of the cytokine mix.

Given these attenuating effects of individual pathway inhibition on cytokine-induced proliferation, we sought to characterize the signaling dynamics of key intracellular signaling pathways in myogenic cells stimulated by the cytokine mix (**Fig. S3A**). We cultured primary myoblasts, the committed myogenic progenitors downstream from MuSCs, on laminin-conjugated 12 kPa hydrogels for 30 min in the presence of the cytokine mix and observed robust activation of phospho-STAT1, phospho-STAT3, phospho-STAT6, and phospho-NFκB by immunoblotting, verifying that the cytokine mix activates its canonical pathways as short term responses (**Fig. S3B-H**). Thus, we hypothesized that similar pathways may be activated in long-term MuSC culture in our platform, and that their temporal activation dynamics may shed light on their roles in long-term MuSC proliferation and differentiation outcomes. MuSCs were cultured on laminin-conjugated 12 kPa hydrogels for five weeks in the presence of the exogenous cytokine mix and lysates were collected weekly for phosphoprotein measurements before passaging (**Fig. 3A**). We used the Luminex multiplexable bead-based platform to measure expression levels of key phosphorylated signaling molecules including phospho-AKT, phospho-ERK1/2, phospho-cJun, phospho-STAT3, phospho-IκBα, phospho-p38α/β, phospho-HSP27, and phospho-STAT1 (all normalized each to β-actin) at each time point. We observed three distinct patterns of phosphoprotein activation dynamics over the long-term culture duration (**Fig. 3B**). First, we detected a delayed biphasic activation signature for phospho-AKT that peaks at 2 wk, returns to baseline at 3 wk, and then continually increases thereafter (**Fig. 3C**). Second, we detected a sustained activation response signature for phospho-ERK1/2, phospho-cJun, phospho-STAT3, and phospho-IκBα that peaks around 1 wk and then attenuates over time (**Fig. 3D**). Third, we detected an oscillatory activation signature for phospho-p38α/β and its downstream effector phospho-HSP27 that peaks at 1 wk, 3 wk and 5 wk (**Fig. 3E**).

**Figure 3.**
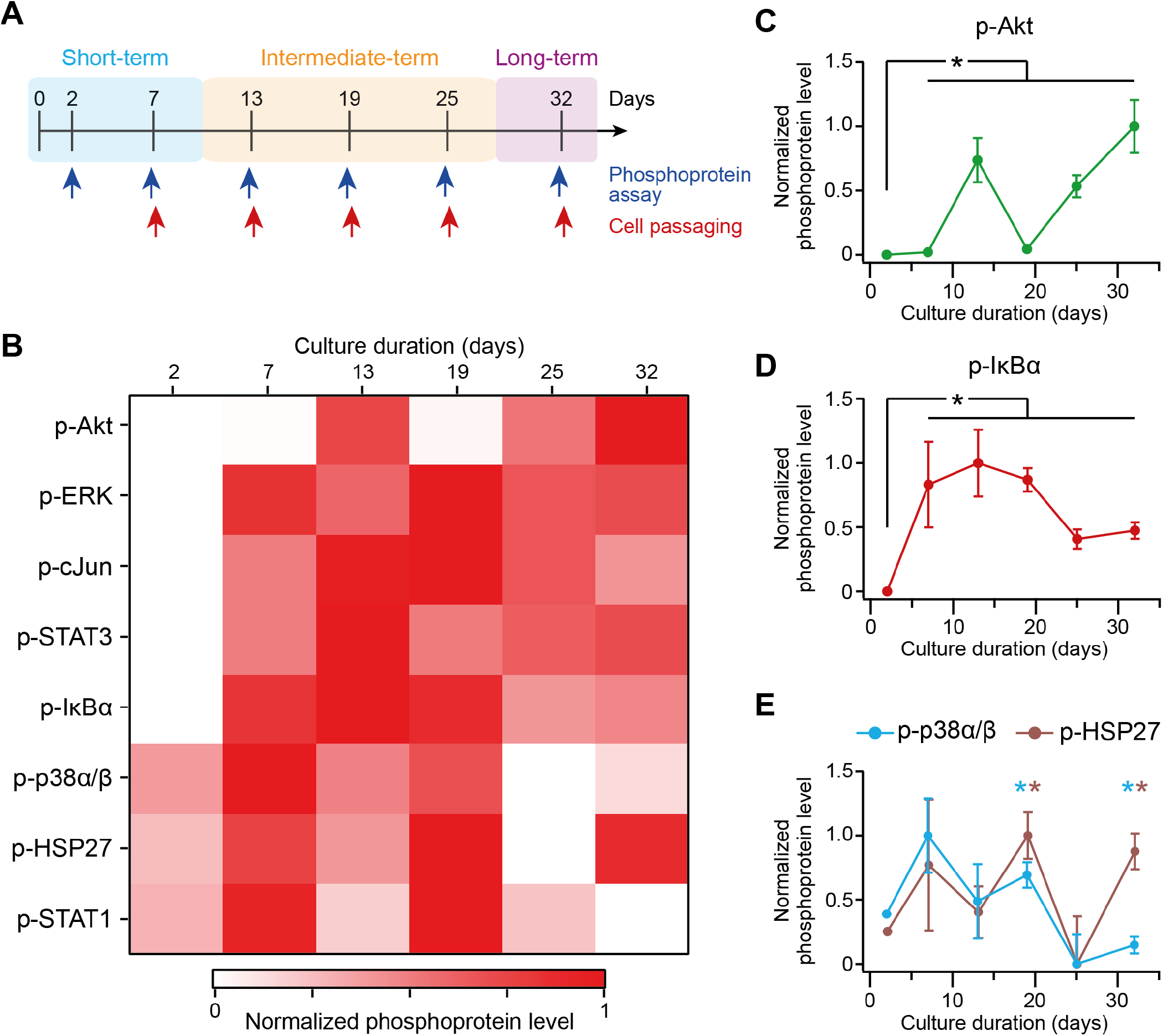
Cytokine mix induces staged intracellular signaling activation during long-term MuSC cultures. (**A**) MuSC were cultured on 12-kPa hydrogels treated with FGF2 and cytokine mix (TNF-α, IL-1α, IL-13, IFN-γ), passaged every 6-8 d, and 8 phosphoproteins were measured using Luminex assays and normalized to β-actin. (**B**-**E**) Normalized phosphoprotein levels at 2-32 d, with each scaled from 0 to 1 over its range. (**B**) Heatmap of mean values, n = 3. (**C**-**E**) Time-courses for p-AKT, p-IκBα, phospho-p38α/β MAPK, and p-HSP27. Mean ± s.e.m., n = 3. * denotes *P* < 0.05 by Student’s T-test compared to d 2. See also **Fig. S3**.

Previous reports have shown that p38α/β induces the activation of quiescent MuSCs (Jones et al., 2005), but also prolonged p38α/β activity contributes to cell cycle exit and induction of myogenic commitment through post-translational regulation of MyoD (Blau et al., 2015; Lluis et al., 2006; Loiben et al., 2017; Perdiguero et al., 2007; Segalés et al., 2016). These distinct functional roles raise the possibility that the initial phase of p38α/β activation within 1 week of culture may be critical for short-term activation and proliferation, whereas p38α/β may promote myogenic differentiation in later phases. We reason that repression of p38α/β-mediated MuSC activation in the first week of culture may explain the diminished the total cellular yield for the p38α/β-inhibited cytokine mix condition (**Fig. 2B**).

### Late-stage inhibition of p38α/β signaling enhances MuSC phenotype in long-term expansion cultures

Based on these p38α/β activation dynamics, we posited that temporally-staged inhibition of p38α/β may improve stem-cell expansion in long-term MuSC cultures on 12-kPa hydrogels stimulated with the cytokine mix. To test this, we cultured adult MuSCs for up to 6 weeks in FGF2 growth medium with or without the cytokine mix and initiated p38i at distinct time points matching each of the passaging events (**Fig. 4A**).

**Figure 4.**
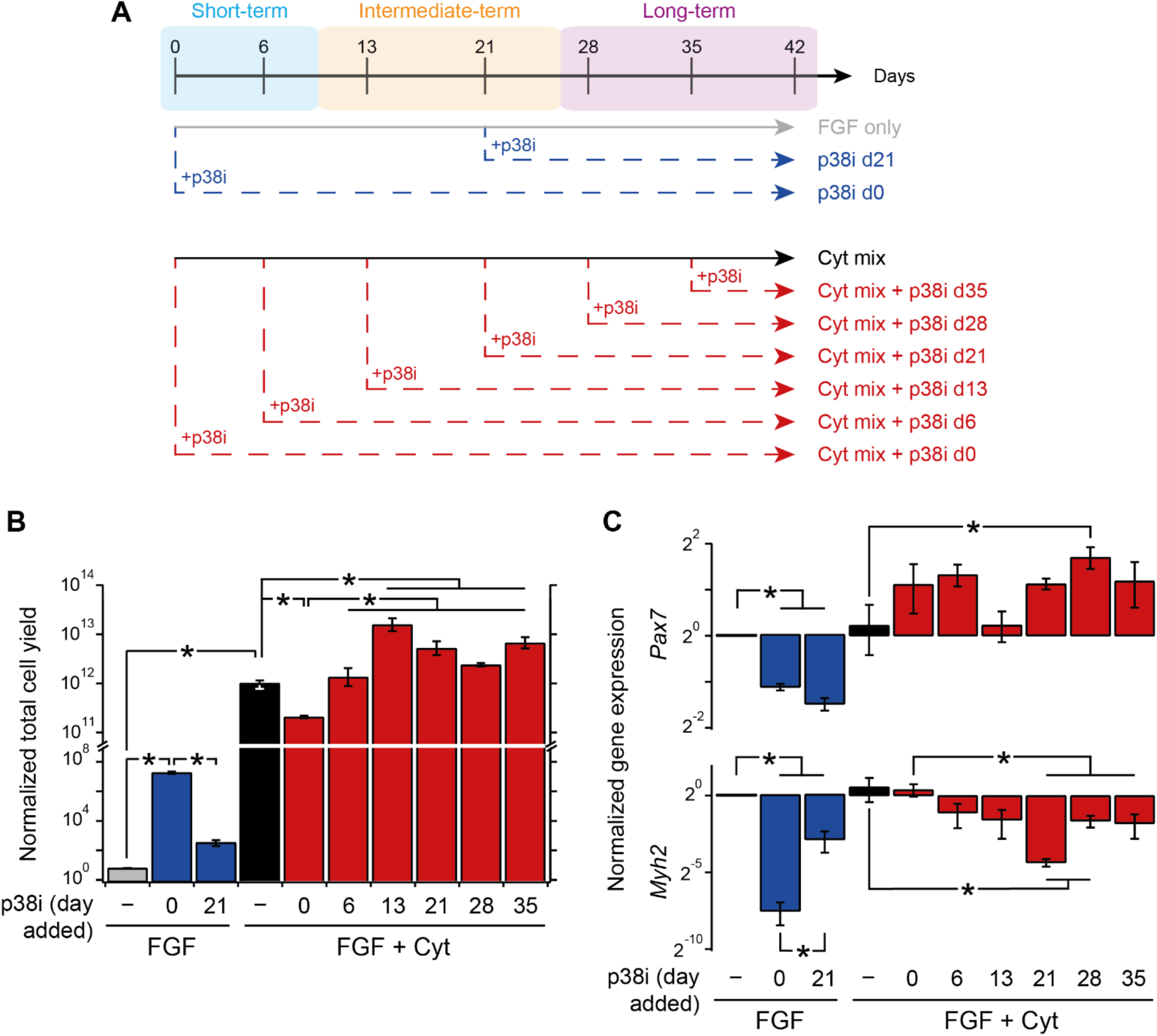
Late-stage inhibition of p38α/β enhances cell yield and stem cell phenotye in long-term MuSC cultures. (**A-C**) Long-term MuSC cultures on 12-kPa laminin-conjugated hydrogels were passaged every 6-8 d and treated with FGF or FGF + cytokine mix (TNF-α, IL-1α, IL-13, IFN-γ), without or with p38i (SB203580, 5 μM; addition staged weekly). (**A**) Schematic of staged p38α/β inhibition scheme. “p38i d*X*” indicates SB203580 was added starting at d *X* and maintained thereafter. (**B**) Normalized cumulative cell yield at 42 d. Mean ± s.e.m., n = 4. See also **Fig. S4** for yield timecourse. (**C**) RT-qPCR assay of *Pax7* and *Myh2* expression normalized to *36B4* at 42 d. Mean ± s.e.m., n = 4. * denotes *P* < 0.05 by Student’s T-test on log-transformed values in (**B**-**C**).

Without cytokine mix, the final total cell yield at 5 weeks was elevated for constant p38i (starting at d 0) relative to no inhibition or when p38i was started at 3 weeks (**Fig. 4B, S4A**). Expression of both the stemness gene *Pax7* and the maturation gene *Myh2* after 6 weeks were diminished by p38i starting at either 0 or 3 weeks, suggesting that long-term p38i in the absence of cytokine mix-stimulation skews MuSC progeny away from a myogenic phenotype (**Fig. 4C**). With cytokine mix, we observed that constant p38i administration suppressed total cell yield by nearly 10-fold and had negligible effects on *Pax7* and *Myh2* gene expression (**Fig. 4B-C**). Notably, this cell yield suppression was not observed for any delayed p38i condition, suggesting that permitting p38α/β activation in the first week of cytokine mix culture improves long-term cell yield. Moreover, delaying the p38i addition until 2 weeks or later of culture enhanced total cell yield above the cytokine mix baseline by a factor of 5-12×. Similar effects were observed in myogenic gene expression; delaying p38i addition until 3-4 weeks of culture enhanced *Pax7* expression and reduced *Myh2* expression, relative to cytokine mix-only controls. These results show that delaying p38α/β inhibition to around 3 weeks of culture enhances MuSC proliferation and stem-cell gene expression while restricting maturation in long-term cytokine mix-stimulated soft hydrogel cultures.

### Enhanced MuSC phenotype with late-stage p38α/β inhibition is contingent on soft hydrogel substrates

To examine the dependency of this enhanced long-term expansion on substrate rigidity, we cultured MuSCs isolated from adult transgenic *Pax7-zsGreen* mice (Bosnakovski et al., 2008) on 12 and 60 kPa laminin-conjugated hydrogels with cytokine mix supplementation, with or without late-stage p38α/β inhibition (from d 21), for five weeks (**Fig. 5A**). MuSCs exposed to late-stage p38i on 12 kPa gels had a distinct, unfused morphology, relative to cells cultured on 60 kPa gels or without p38i (Fig. 5B). In both 12 and 60 kPa conditions, p38α/β inhibition significantly increased the cumulative cell yield at five weeks, but there was no difference in total yield between any 12 and 60 kPa conditions (**Fig. 5C-D**).

**Figure 5.**
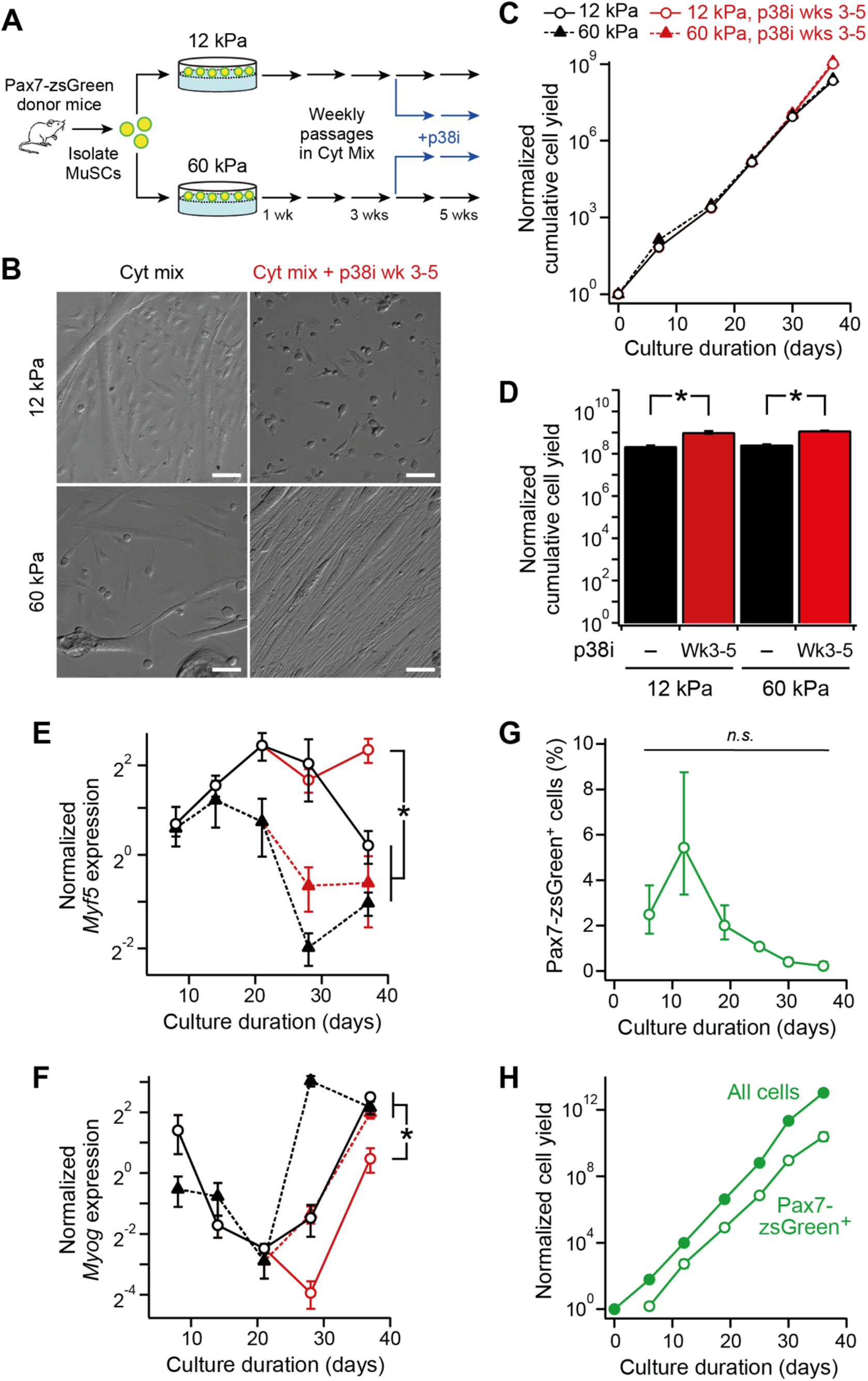
Late-stage p38α/β inhibition enhancement of MuSC phenotypes are contingent on soft hydrogel substrates. (**A-H**) MuSC isolated from *Pax7-zsGreen* transgenic mice were cultured 12 or 60 kPa laminin-conjugated hydrogels were passaged every 7 d and treated with FGF and cytokine mix (TNF-α, IL-1α, IL-13, IFN-γ), with or without p38i (SB203580, 5 μM) from 3 to 5 wks. (**A**) Experimental scheme. (**B**) Representative modulation contrast images. Scale bar, 100 μm. (**C**) Normalized cumulative cell yield from 0-5 wks. Mean ± s.e.m., n = 3. (**D**) Normalized total cell yield at 5 wks. Mean ± s.e.m., n = 3. (**E-F**) RT-qPCR measurement of *Myf5* and *Myog* expression normalized to *36B4* at 1-5 wks. Mean ± s.e.m., n = 4. (**G**) Normalized cumulative and Pax7-zsGreen^+^ cell yield for cytokine mix with p38i for 3-5 wks on 12 kPa hydrogels. Mean ± s.e.m., n = 3. (**H**) Pax7-zsGreen^+^ cells percentage for cytokine mix with p38i for 3-5 wks on 12 kPa hydrogels. Mean ± s.e.m., n = 3. * denotes *P* < 0.05 by Student’s t-test or not significant (n.s.) on log-transformed values in (**C**-**F**) and on non-transformed values in (**G**).

We performed RT-qPCR at each passage throughout the long-term culture to examine the dynamics of myogenic differentiation. We found that *Myf5*, a marker of MuSC activation, remained elevated and that *Myog*, a myogenic commitment marker, remained suppressed only in the 12 kPa late-stage p38i condition (**Fig. 5E-F**), suggesting the soft but not stiff substrates with late-stage p38α/β inhibition sustain an activated MuSC phenotype.

For the late-stage p38i 12-kPa condition, we analyzed zsGreen transgene expression by live-cell microscopy as a more direct measure of Pax7-expressing MuSC cells (**Fig. 5G-H**). The frequency of Pax7-zsGreen^+^ cells ranged between ~0.2-5.4% throughout the long-term culture but specific differences between timepoints were not statistically significant (**Fig. 5G**). By comparing to the extrapolated total cell yield, we estimate that this protocol yields ~10^9^-10^10^ Pax7-zsGreen^+^ cells by 5 weeks from each sorted and seeded MuSC at d 0 (**Fig. 5G**). These results suggest the soft (12 kPa) hydrogel substrate with late-stage p38α/β inhibition permits robust and prolonged expansion of an activated MuSC population, whereas the stiffer (60 kPa) substrate is able to support similar yields but with a more committed myogenic cell phenotype.

### Cytokine mix with late-stage p38α/β inhibition maintains MuSC engraftment potential in long-term culture

Pax7 expression status is a hallmark of the stem cell phenotype of MuSCs, but confirmation of MuSC functionality requires *in vivo* transplantation assay. To determine whether the long-term culture protocol maintains functional MuSCs, we performed a sensitive *in vivo* transplantation engraftment assay (Cosgrove et al., 2014; Gilbert et al., 2010; Sacco et al., 2008) on MuSCs during long-term cultures. We isolated MuSCs from adult *Luciferase* transgenic donor mice and cultured them on 12 or 60 kPa laminin-conjugated hydrogels with cytokine mix supplementation and with or without late-stage (weeks 3-5) p38α/β inhibition. We collected cells at 3 or 5 weeks of culture and transplanted 500 cells into immunocompromised *NSG* recipient mice (**Fig. 6A**). One month post-transplantation, we measured *in vivo* cell engraftment using a bioluminescent imaging (BLI) assay that is sensitive to individual myofiber engraftment events (Cosgrove et al., 2014; Sacco et al., 2008). Mice transplanted with MuSCs cultured on 12 kPa hydrogels with cytokine mix for 3 wk exhibited substantial engraftment (67% of transplants), but cells from 5 wk did not (0%), suggesting that functional MuSCs were no longer present at the end of the culture protocol (**Fig. 6B**). In contrast, if p38α/β was inhibited from 3-5 wk under this condition, we observed robust engraftment outcomes at similar frequencies (100%) and levels as from 3 wk of cytokine mix culture, suggesting that late-stage p38i extends the preservation of MuSCs function to 5 wk. We observed similar results for MuSCs cultured on the stiffer 60 kPa hydrogels, though the engraftment levels were reduced relative to the 12 kPa conditions. Critically, we observed negligible engraftment frequencies for MuSCs cultured on traditional laminin-coated tissue-culture polystyrene substrates (~3 GPa Young’s modulus (Carraher Jr., 2016)) with cytokine mix treatment, regardless of p38α/β inhibition (**Fig. S5A-B**). These results indicate soft substrates (~12 kPa) permit maintenance of a stem cell-like engraftment potential for up to 5 weeks of culture in the presence of the cytokine mix and late-stage p38α/β inhibition, but more rigid substrates are not permissive to this preservation of functional MuSC potential during long-term cultures.

**Figure 6.**
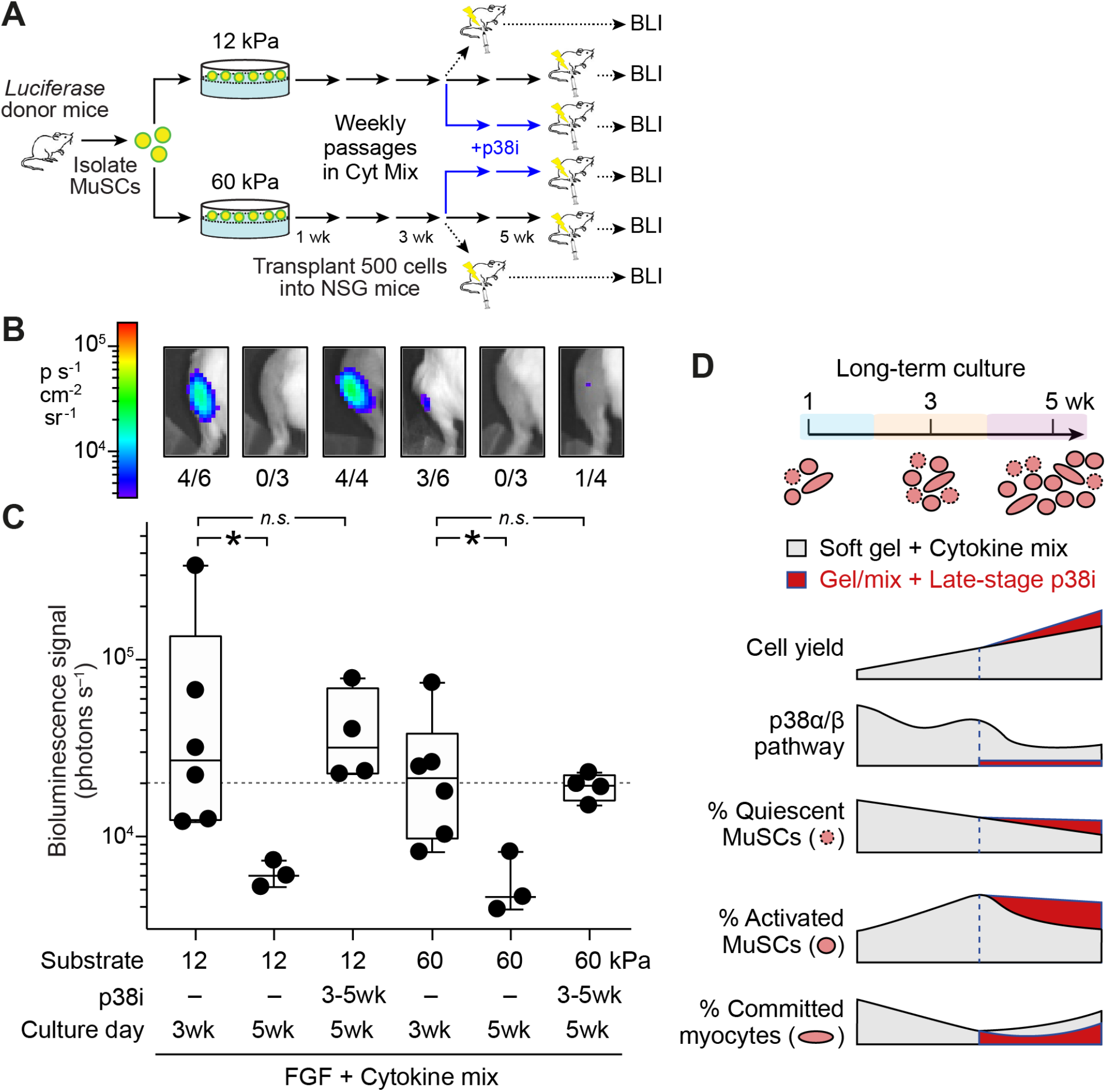
Late-stage inhibition of p38α/β MAPK signaling enhances MuSC engraftment in *in vivo* transplantation. Luciferase-expressing MuSCs were cultured on 12 or 60 kPa laminin-coated hydrogels in the presence of cytokine mix mix (TNF-α, IL-1α, IL-13, IFN-γ) with or without the addition of p38i starting from 3-5 wks. At 3 wk or 5 wk, 500 cells were transplanted into NSG mice and engraftment was measured using bioluminescent imaging (BLI) 1-month post-transplantation. (**A**) Transplant assay schematic. (**B**-**C**) Background-subtracted representative BLI images (**B**) and signals (**C**) at one-month post-transplantation. Dashed line indicates positive engraftment threshold of 20,000 photons s^−1^. * denotes *P* < 0.05 or not significant (n.s.) by Mann-Whitney U test. Fraction of transplants resulting in positive engraftment outcome reported in inset of (**B**). See also **Fig. S5.** (**D**) Summary schematic.

## Discussion

This work presents an advance in long-term *ex vivo* MuSC expansion methods. We demonstrated that a TNF-α/IL-1α/IL-13/IFN-γ cytokine mix supplement to standard FGF2-containing myogenic growth medium supports the proliferative expansion of muscle stem cells *ex vivo,* which are capable potent engraftment function *in vivo*, and these outcomes are optimized on soft, muscle-mimicking (~12 kPa) culture substrates in combination with a delayed inhibition of the p38α/β MAPK signaling pathway (**Fig. 6C**). Together, our data suggest that the expanded cells under the late-stage p38i condition contain a heterogenous population of Pax7^+^ stem cells (~1%, using a sensitive Pax7-zsGreen reporter system) and activated stem/progenitor cells, with infrequent committed or fused cells (**Fig. 5**). Intriguingly, we observed enhanced transplantation potential for MuSCs maintained on 12 kPa laminin-conjugated hydrogels compared to laminin coated plastic, suggesting substrate rigidity can modulate long-term engraftment potential of MuSCs, as has been reported previously for short-term cultures (Gilbert et al., 2010). In our short-term studies, we did not observe substantial differences in MuSC phenotype between 5-60 kPa substrates, though the bulk MuSC phenotype assays reported in **Fig. 1** may not sufficiently distinguish between the early self-renewal division differences within the 2-40 kPa range that were reported by Gilbert et al.

Our findings suggest that chronic administration of the TNF-α/IL-1α/IL-13/IFN-γ cytokine mix is a potent driver of exponential MuSC proliferation for at least one month of culture, in agreement with Fu et al. (Fu et al., 2015). These cytokines are not chronically present in homeostatic muscles, and instead are dynamically regulated with the transient cycles of T-cell infiltration during healthy muscle regeneration processes (De Micheli et al., 2020; Tidball, 2017). The prolonged cytokine stimulation protocol tested herein drives a diverse set of signaling network activation signatures, including phosphoprotein pathways with delayed, biphasic, or oscillatory dynamics (**Fig. 3**). Most of these cytokine-induced signaling pathways provide critical pro-proliferative contributions in the initial phase of MuSC proliferation (**Fig. S2C**), and may also be responsible for the observed shift towards a Myosin Heavy Chain-expressing population of committed myocytes and myotubes in the late culture stages (**Figs. 4C** and **5F**).

Notably, we found that the p38α/β pathway exhibited a prolonged oscillatory activation signature and that a delayed pharmacological inhibition of p38α/β ameliorated its pro-differentiation effects and permitted MuSC function in long-term cultures. These findings are in agreement with prior reports in short-term culture models (Bernet et al., 2014; Charville et al., 2015; Cosgrove et al., 2014; Jones et al., 2005; Palacios et al., 2010) and p38α-specific genetic ablation studies (Brien et al., 2013) demonstrating that the p38α/β pathway drives both myogenic proliferation and differentiation. Further, our observations suggest that p38α/β exhibits temporally disparate effects in the regulation of MuSC self-renewal and differentiation under this cytokine mix long-term culture system. We have previously observed, through data-modeling of myoblast cell fates, that the p38α/β pathway exhibits time-varying effects on myogenic proliferation and commitment in short-term (~24 hr) stimulation periods (Loiben et al., 2017). These findings further suggest that p38α/β pathway dynamics may encode differing consequences for myogenic outcomes; thus, more time-resolved inhibition strategies may provide further enhancement of MuSC expansion outcomes. Similarly, combinatorial targeting of other signaling pathways, such as through inhibition of STAT3 signaling (Price et al., 2014; Tierney et al., 2014), may further support MuSC self-renewal in long-term cytokine mix cultures.

Likewise, this protocol may be further enhanced through strategies to promote a shift toward a more quiescent MuSC population before transplantation. Recently, Sampath et al. have reported oncostatin-M (OSM) promotes MuSC quiescence through cell-cycle exit and enhances their serial transplantation potential (Sampath et al., 2018). Addition of OSM or other pro-quiescence strategies (Quarta et al., 2016), possibly timed in a manner that better mimics the *in vivo* dynamics of the post-injury return to homeostasis, may provide a synergistic improvement in MuSC functional potency in long-term expansion cultures. Moreover, *ex vivo* expansion strategies such as presented here could be combined with cell delivery technologies consisting of synthetic and/or natural material (Davoudi et al., 2018; Han et al., 2018; Rao et al., 2017; Sleep et al., 2017; Wolf et al., 2015) to improve cell-engraftment outcomes and functional recovery endpoints. The advances presented in this work provide an *ex vivo* MuSC expansion protocol capable of reaching the high-yield functional expansion demands for clinical muscle cell therapy applications. Further optimization of this technology and its extension to human muscle stem cells from healthy and diseased patients could help realize MuSC-based therapies for chronic muscle diseases.

## Materials and Methods

### Mice

The Cornell University Institutional Animal Care and Use Committee (IACUC) approved all animal protocols and experiments were performed in compliance with its institutional guidelines. Mouse housing and husbandry was conducted in vivaria managed by the Cornell Center on Animal Research and Education or as service from the Cornell Progressive Assessment of Therapeutics (PATh) Facility (for NSG mice). C57BL/6J wildtype and dystrophic *mdx* (C57BL/10ScSn-*Dmd^mdx^*/J) mice were obtained from Jackson Laboratory (# 000664 and # 001801, respectively). *Pax7-zsGreen* transgenic mice (Bosnakovski et al., 2008) were a generous gift from M. Kyba (University of Minnesota). These strains were maintained heterozygously on a C57BL/6J background, routinely genotyped, and bred in-house. Transplant experiments used transgenic *Luciferase* mice (FVB-Tg(CAG-luc,-GFP)L2G85Chco/J; Jackson Laboratory # 008450) as donors and immunodeficient NOD/Scid/IL2Rg^null^ (NSG) (NOD.Cg-*Prkdc^scid^ Il2rg^tm1Wjl^*/SzJ; Jackson Laboratory # 005557) as recipients. All experiments used adult 3-5 month-old mice. A random mixture of male and female mice was used in each experiment, except for the mdx studies for which only male mice were used, with n = 3-4 donor mice pooled for each MuSC isolation.

### MuSC isolation

MuSCs were isolated by FACS from the hindlimbs muscles of adult mice following established protocols (Sacco et al., 2008). In brief, following dissociation and magnetic depletion, MuSCs were prospectively isolated using a (propidium iodide/CD45/CD11b/CD31/Sca1)^−^ CD34^+^ α7-integrin^+^ cell sorting gate. In detail, we euthanized mice with isoflurane and harvested tibialis anterior, quadriceps, and gastrocnemius muscles. Muscles were digested with 2.5 mg ml^−1^ Collagenase D (Sigma-Aldrich # 11088866001) and 0.04 U ml^−1^ Dispase II (Sigma-Aldrich # 04942078001) followed by dissociation using a gentleMACS system (Miltenyi Biotec # 130-093-235). Cell suspensions were filtered through 100 and 40 μm filters (Corning Cellgro # 431752 and #431750) to remove myofiber debris. Erythrocytes were removed through incubation in erythrocyte lysis buffer (IBI Scientific # 89135-030) supplemented with 0.1% DNAse (Omega Bio-Tek # E1091-02). Cell suspensions were stained with biotinylated antibodies against CD45 (Biolegend # 103104), CD31 (Biolegend # 102404), CD11b (Biolegend # 101204), and Sca1 (Biolegend # 108104) for 15 min at 4°C, washed, and then stained with streptavidin-conjugated magnetic beads (Miltenyi Biotec # 130-048-102), streptavidin–PE/Cy7 (Biolegend # 405206), and antibodies against α7-integrin (Alexa Fluor 647; AbLab # 67-0010-05), CD34 (eFluor 450; Thermo Fisher Scientific # 48-0341-82) for 20 min at 4°C. (CD45/CD11b/CD31/Sca1)-positive cells were depleted by passage through LS selection columns (Miltenyi Biotec # 130-042-401). All washes, staining steps, and resuspension for FACS was performed in a FACS buffer solution containing 5% goat serum (Jackson Immunoresearch #005-000-121) and 1 mM EDTA in 1× phosphate buffer saline (PBS). Suspensions were stained with propidium iodide (PI, Thermo Fisher Scientific # P3566) immediately prior to FACS. FACS was performed on an FACSAria Fusion sorter (BD Biosciences).

### Hydrogel fabrication

Engineered 2D basal lamina constructs were developed using previously established methods (Cosgrove et al., 2014; Gilbert et al., 2010; Lutolf and Hubbell, 2003). 10 kDa molecular weight 4-arm polyethylene glycol (PEG)-thiol (PEG-VS; JenKem # A7008-1) and 10 kDa 8-arm PEG-vinyl sulfone (PEG-SH; JenKem # A10033-1) were dissolved triethanolamine and DI water, respectively, at 10% wt/vol. Precleaned glass slides were coated with a thin layer of Sigmacote (Sigma-Aldrich, # SL2-100ML) and baked for 4 h at 80°C. PEG-SH and PEG-VS were briefly mixed at a 2:1 ratio in sufficient TEOA to match desired PEG wt% and 150 uL aliquots were pipetted onto a baked glass slide, with another slide placed on top of two 0.8 mm plastic spacers. The whole assembly was secured with binder clips and incubated for 12 min at 37°C. The setup was then disassembled, and the solidified gels were each coated with 30 μL of a pre-dialyzed laminin (0.125 μg μL^−1^ in 1× PBS; Thermo Fisher Scientific, # 23017015). The gels were incubated for 50 min to complete the laminin conjugation reaction and then were attached to the bottoms of 24-well tissue-culture plastic plates with 10 μL of the PEG-SH/PEG-VS/TEOA mixture per gel. Gels were stored at 4°C in PBS with 1% antibiotic-antimycotic (Corning Cellgro # 30-004-CI) for up to 1 wk. Gels were washed with cell culture medium 3× prior to cell seeding.

### Rheological testing of hydrogels

Rheological testing was performed at the Cornell Energy Systems Institute using an Anton Paar MCR rheometer (both 301 and 501 models). For ease of handling and testing, the machine was configured to perform rheology using 10-mm diameter parallel plate settings. Hydrogels were kept in PBS with 1% antibiotic-antimycotic prior to testing to avoid dehydration. Shear rheometry was performed with minimal compression (≤0.1 N) to achieve no-slip shear measurements at 5% angular shear strain percentage and a 60–0.6 Hz oscillation frequency range with 8 points per decade analyzed in the frequency range. Storage (G’) and loss (G’’) modulus values were calculated and converted to a Young’s modulus (E) values for hydrogels following E = 2G(1+**v**) where G is the shear storage modulus at 6 Hz and **v** represents the hydrogel’s Poisson ratio, estimated to be 0.5. The mean Young’s modulus from n = 3 independent replicates was related to the PEG weight percentage (**Fig. S1A**) by fitting to a 4-PL model, which was used to identify PEG weight percentages needed to achieve targeted moduli.

### Protein incorporation assays of hydrogels

PEG hydrogels were synthesized at 3.0, 4.0, or 5.0 wt% PEG and were conjugated as described above with 4 μg cm^−2^ of dialyzed laminin in 1× PBS. Gels were incubated with cysteamine (Sigma-Aldrich # 30070-10G) for 1 h to terminate unreacted PEG-VS arms and then rinsed with 0.05% Tween-20 in 1× Tris-buffered saline (TBS-Tween). Hydrogels were blocked with 5% bovine serum albumin (BSA) in TBS-Tween for 2 h, then washed twice with TBS-Tween. Hydrogels were then incubated with an anti-laminin antibody (1:200 dilution in 1% BSA; Sigma-Aldrich # L9393) for 1 h, then washed five times with TBS-Tween. Wells were then incubated with peroxidase-conjugated goat anti-rabbit secondary antibody (1:500 dilution in 1% BSA; Jackson Immunoresearch # 111-035-144) for 1 h, then washed five times with TBS-Tween. Hydrogels were incubated in ECL reagent (1:1 mix of luminol/HRP substrate solutions; Bio-Rad # 1705062) for 1 min and then imaged for 300 s using the ChemiDoc imaging system (Bio-Rad # 17001401). Dialyzed laminin was adsorbed directly to wells to generate a standard reference curve for quantitation. Luminescence images were analyzed using ImageLab (Bio-Rad) software.

### Short-term MuSC culture

Isolated MuSCs were seeded at 1000 cells cm^−2^ on laminin-conjugated 5, 12, 20, 30, or 60 kPa Young’s modulus PEG hydrogels in 24-well plates. MuSCs were cultured in 2 mL myogenic growth medium (GM) containing 43% Dulbecco’s Modified Eagle’s Medium (DMEM; Corning Cellgro # 10-013), 40% Ham’s F-10 (Corning Cellgro # 10-070-CV), 15% fetal bovine serum (Corning Cellgro # 35-010-CV), 1% Penicillin-Streptomycin (Corning Cellgro # 30-002-CI), 1% L-glutamine (Corning Cellgro # 25-005), and 2.5 ng mL^−1^ recombinant mouse FGF2 (R&D Systems # 3139-FB-025). In some experiments, a mix of recombinant mouse cytokines: 10 ng mL^−1^ TNF-α (R&D Systems # 410-MT-010), 10 ng mL^−1^ IL-1α (R&D Systems # 400-ML-005), 10 ng mL^−1^ IL-13 (R&D Systems # 413-ML-005), and 10 ng mL^−1^ IFN-γ (R&D Systems # 485-MI-100) was added. For pathway inhibition studies, the following chemicals were added: 5 μM SB203580 (p38α/β MAPK inhibitor; Selleck Chemicals # S1076), 10 nM PD0325901 (MEK1/2 inhibitor; Selleck Chemicals # S1036), 1 μM SP600125 (pan-JNK inhibitor; Selleck Chemicals # S1460), 5 μM 5,15-diphenyl-porphine (STAT3 inhibitor; Sigma-Aldrich # D4071), 10 nM GSK2110183 (AKT1/2/3 inhibitor; Selleck Chemicals #S7521), 1 μM AG-490 (JAK2 inhibitor with effects on EGFR kinase; Selleck Chemicals # S1143), or 10 μM BMS-345541 (IKK-1/2 inhibitor; Selleck Chemicals # S8044), all resuspended in 0.1% DMSO final concentration. Media was replenished every 2 d. At 7 d, cells were lifted from gels using 0.25% Trypsin/0.1% EDTA (Corning Cellgro # 25-053-CI) and quenched with GM. Samples were counted using a hemocytometer or lysed for RNA isolation.

### Long-term MuSC culture

Isolated MuSCs were seeded at 1000 cells cm^−2^ on laminin-conjugated 12 or 60 kPa PEG hydrogels in 24-well plates. MuSCs were cultured in 2 mL myogenic growth medium (GM) with FGF2 as described above. For some experiments, a mix of recombinant mouse cytokines (10 ng mL^−1^ TNF-α, 10 ng mL^−1^ IL-1α, 10 ng mL^−1^ IL-13, 10 ng mL^−1^ IFN-γ) and/or 5 μM SB203580 (p38α/β MAPK inhibitor) were added. Media were changed every 3 d. Every 6-8 d, cells were passaged by lifting from the gels using 0.25% Trypsin/0.1% EDTA, quenched with GM, and counted with a hemocytometer. After counting, cells were pooled in equal numbers from n = 1-4 wells per condition and were reseeded at 500-1000 cells cm^−2^ on new laminin-conjugated hydrogels and in fresh medium. In some experiments, wells were fixed or lysed for other analyses. Long-term cultures were maintained by weekly passages until 28-42 d. For experiments on plastic substrate in **Fig. S5**, 24-well tissue-culture plastic plates were coated with 50 μL dialyzed laminin (0.125 μg μL^−1^ in 1× PBS), incubated at 37°C, and rinsed three times with 1× PBS prior to seeding.

### Modulation contrast microscopy

Cells were washed 3× with cold 1× PBS, then incubated with cold 4% paraformaldehyde in PBS for 12 min. Cells were washed 3× with cold 1× PBS, then left in 1× PBS for imaging. Modulation contrast images were acquired using a modified Nikon Eclipse Ti-E microscope (Micro-Video Instruments, Inc.) with custom green LED light source, a Nikon LWD NAMC 20× objective (# MRP66205), and an Andor Zyla 5.5 scMOS Camera. Digital images were captured with 50 ms exposure.

### Quantitative immunoblotting

Primary myoblasts (PMBs) were isolated from C57BL/6J mice as described previously (Rando and Blau, 1994). PMBs were seeded on 12 kPa PEG hydrogels in GM with FGF2, cultured for 12-18 h, then switched to DMEM without serum for 6 h, and stimulated with or without the cytokine mix for 30 min. Cells were washed with cold 1× PBS and lysed for immunoblotting using an NP-40-based lysis buffer containing 50 mM b-glycerophosphate, 30 mM NaF, 10 mM NaPP, 50 mM Tris– HCl, 0.5% NP-40 substitute, 150 mM NaCl, 1 mM benzamidine, 2 mM EGTA, 400 μM sodium orthovanadate, 200 μM DTT, 2 mM PMSF, 1:200 dilution Phosphatase Inhibitor Cocktail Set III (EMD Millipore # 524627). Lysate were collected and centrifuged at 4°C for 10 min at 15,000×*g*. The lysate supernatants were collected, and protein concentration was quantified using a Micro BCA Protein Assay Kit (Thermo Fisher Scientific # 23235) per manufacturer’s protocol. Electrophoresis gels (1.5 mm thickness, 10% acryl/bisacrylamide, Tris-HCl, ammonium persulfate, tetramethylethylenediamine, sodium dodecyl sulfate (SDS)) were loaded with 25 μg of sample in 25 μL of 1× sample buffer (20 mM Tris-HCl, Glycine, 10% SDS, 0.4% β-mercaptoethanol) per lane or 1 μL of strep-tagged unstained protein standards (Bio-Rad # 1610363) and run at 100 V for 2 h in Tris-HCl-Glycine-SDS running buffer. Proteins were transferred to a methanol-activated PVDF membrane overnight at 4°C, 15 V in Tris-HCL/Glycine/Methanol transfer buffer. Membranes were blocked in 5% powdered milk in Tris-buffered saline with Tween-20 (TBST) with gentle rocking at room temperature for 1 hr. Primary antibodies (anti-phospho-STAT1, anti-phospho-STAT3, anti-phospho-STAT6, anti-phospho-NFκB, anti-GAPDH, or anti-HSP90; see table for details) were diluted in 5% powdered milk in TBST and blots were incubated with diluted antibodies with gentle rocking at room temperature for 1 hr. Blots were then washed three times with 1× TBST for 5 min per wash. Peroxidase-conjugated goat anti-rabbit (Jackson Immunoresearch # 111-035-144) and/or peroxidase-conjugated goat anti-mouse (Jackson Immunoresearch # 115-035-146) secondary antibodies were diluted 1:200 in 1× TBST and incubated with blots under gentle rocking at room temperature for 30 min. Blots were then washed three times with 1× TBST for 5 min per wash. Blots were incubated in ECL reagents (1:1 mix of luminol/HRP substrate solutions; Bio-Rad # 1705062) for 1 min and then imaged for 120 s using the ChemiDoc imaging system (Bio-Rad # 17001401). Blots were analyzed using ImageLab software (BioRad) to calculate individual band intensities.

| Target | MW (kDa) | Type | Dilution | Vendor | Catalog # |
| --- | --- | --- | --- | --- | --- |
| p-STAT1 (pY701) | 85 | Monoclonal rabbit | 1:1000 | Cell Signaling Technology | 9167S |
| p-STAT3 (pY705) | 80 | Monoclonal rabbit | 1:1000 | Cell Signaling Technology | 9145S |
| p-STAT6 (pY641) | 110 | Monoclonal rabbit | 1:1000 | Cell Signaling Technology | 56554S |
| p-NFκB (pS536) | 65 | Monoclonal rabbit | 1:1000 | Cell Signaling Technology | 3033S |
| GAPDH | 37 | Monoclonal mouse | 1:2500 | Thermo Fisher Scientific | AM4300 |
| HSP90 | 90 | Polyclonal rabbit | 1:2500 | Santa Cruz Biotechnology | sc-7947 |

### Quantitative RT-PCR

We isolated RNA from cell pellets using the E.Z.N.A. MicroElute Total RNA Kit (Omega Bio-tek # R6831-01) into 30 μL of protease-free water. cDNA was obtained via reverse transcription using the High Capacity cDNA RT Kit (Thermo Fisher Scientific # 4368814) and prepared for RT-PCR using the SYBR Green PCR MasterMix (Thermo Fisher Scientific # 4309155). PCR was performed in a Viia 7 Real-Time PCR System (Thermo Fisher Scientific) using the following settings: cycling at 95°C for 10 min followed by 40 cycles of 95°C for 15 sec and 60 °C for 1 min. We quantified transcript levels using the 2^−ΔΔCT^ method to compare gene expression levels between treatment conditions and appropriate controls. Primer sequences were used for *Pax7*, *Myf5, Myod1*, *Myog*, and *36B4* are reported in the table below).

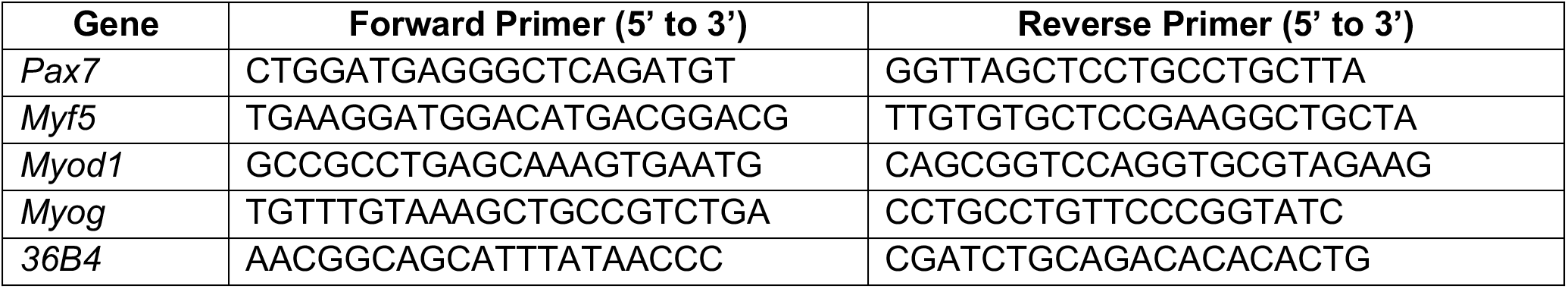

### Luminex phosphoprotein immunoassays

MuSCs were cultured on 12 kPa laminin-coated hydrogels as described above. As part of the long-term MuSC culture passaging protocol, cells were lysed (n = 3-4 replicates per condition) on days 2, 7, 13, 19, 25, and 32. Cells were washed with cold 1× PBS and lysed using the Bio-Plex Pro Cell Signaling Reagent Kit lysis buffer (Bio Rad # 171304006M) supplemented with 2 mM PMSF and 1:200 dilution Phosphatase Inhibitor Cocktail Set III (EMD Millipore # 524627). Lysates were collected and centrifuged at 4°C for 10 min at 15,000×*g*. Supernatants were collected and protein concentrations were quantified using a Micro BCA Protein Assay Kit per manufacturer’s protocol. Bio-Plex Pro Magnetic Cell Signaling assays (Bio Rad) were used to quantify the following phosphoproteins: phospho-AKT (Ser473; # 171V50001M), phospho-ERK1/2 (Thr202/Tyr204, Thr185/Tyr187; # 171V50006M), phospho-cJun (Ser463; 171V50003M), phospho-STAT3 (Tyr705; # 171V50022M), phospho-IκBα (Ser32/Ser36; # 171V50010M), phospho-p38α/β MAPK (Thr180/Tyr182; # 171V50014M), and phospho-HSP27 (Ser78; # 171V50029M). Assays were performed in multiplex using the Bio-Plex Pro Cell Signaling Reagent Kit per the manufacturer’s protocol with 5 μg of sample per assay, n = 1 technical replicate and n = 3-4 biological replicates per time point. Background-subtracted fluorescence values for each phosphoprotein were normalized to the background-subtracted fluorescence values for Bio-Plex Pro Magnetic Signaling Assay Total β-Actin (Bio Rad # 171V60020M). Normalized values were averaged across biological replicates and then scaled from 0 to 1, with 0 corresponding with the lowest and 1 corresponding with the highest value across time points for a given phosphoprotein.

### Imaging *Pax7-zsGreen* transgene expression

MuSCs isolated by FACS from Pax7-zsGreen transgenic mice by FACS using (propidium iodide/CD45/CD11b/CD31/Sca1)^−^ CD34^+^ α7-integrin^+^ cell sorting gate (typically with >95% zsGreen positivity; data not shown) and were maintain in long-term cultures as described above. Modulation contrast and epifluorescence images were acquired using a modified Nikon Eclipse Ti-E microscope (Micro-Video Instruments, Inc.) with a custom green LED light source, a 470-nm excitation source from a SPECTRA-X light engine (Lumencor), a Nikon LWD NAMC 20× objective (# MRP66205), a DAPI/FITC/Cy3/Cy5 polychroic (Chroma # VCGR-SPX-P01), and an Andor Zyla 5.5 scMOS Camera. Modulation contrast and fluorescence images were captured with 50 ms and 1 s exposures, respectively. Cell Profiler (ver. 3.0.0, Broad Institute) was used to segment and threshold images. Median intensity of zsGreen signal for each segmented cell was calculated and then background subtracted to normalize intensity across each day/condition. For each culture day, a consistent threshold was applied to background-subtracted median intensities to assign a positive or negative Pax7-zsGreen expression state to each cell.

### MuSC transplantation engraftment assays

Cell transplantation assays were performed to assess the engraftment potential of cultured MuSCs, following previous reports (Cosgrove et al., 2014; Gilbert et al., 2010; Quarta et al., 2016; Sacco et al., 2008). MuSCs were isolated from transgenic Luciferase mice, with Luciferase expression regulated by the ubiquitous CAG promoter. After specific culture durations, cells were collected, counted, and resuspended into FACS buffer solution. Transplants were performed with 500 cells per 10 μL into the tibialis anterior muscles of anesthetized recipient NSG mice by intramuscular injection. Bioluminescent imaging (BLI) was performed at 1 mo post-transplant to assess transplanted cell survival and engraftment. Recipient mice were anesthetized with isoflurane, administered 0.1 mmol kg^−1^ D-luciferin reconstituted in 150 μL sterile 1× PBS by intraperitoneal injection, and imaged on an IVIS Spectrum In Vivo Imaging System (Perkin Elmer) 12 min later. BLI images were analyzed using Living Image (Perkin Elmer) software with a fixed region-of-interest (ROI) size to quantify the bioluminescent signal from each hindlimb. BLI thresholds indicated for stable cell engraftment were set (in **Figs. 6C** and **S5B**) in agreement with previous reports (Cosgrove et al., 2014; Sleep et al., 2017).

### Statistical analysis

All cell culture experiments were performed with n = 3-4 replicates unless otherwise noted. Transplant engraftment assays were performed using n = 3-10 replicates and BLI levels were analyzed using a Mann-Whitney U test. An unpaired two-tailed (Student’s) T test was used for all other data. Notably, cumulative cell yield and relative gene expression values were both log-transformed prior to statistical testing. A significance level α = 0.05 was used for all statistical tests. In figures, **\*** denotes a p-value < α and *n.s.* (not significant) denotes a p-value ≥ α.

## Acknowledgments

This work was financially supported by the National Institutes of Health under awards R00AG042491, R01AG058630, R21EB024747, and R21AR072265 (to B.D.C), the National Science Foundation under award DMR-1006323 (to L.A.A.), a US Department of Education Graduate Assistantship in Areas of National Need under Award P200A150273 (to A.M.L.), Roberta G. and John B. DeVries Graduate Fellowships (to A.M.L. and V.M.A.), Hunter R. Rawlings III Cornell Presidential Research Scholarships (to J.C.C. and R.F.K.), and a Cornell Engineering Learning Initiatives Undergraduate Research Award (to P.F.). The authors acknowledge technical support from the Cornell Center for Animal Resources and Education and Cornell University Biotechnology Resource Center (BRC) Flow Cytometry Facility and Imaging Facility. The IVIS-Spectrum imaging system was supported by NIH award S10OD025049. The authors acknowledge support on mechanical rheometry instrumentation from the Cornell Energy Systems Institute (CESI). The authors acknowledge the kind donation of Pax7-zsGreen transgenic mice from M. Kyba at the University of Minnesota. The authors acknowledge technical assistance from A. De Micheli, G. Livermore, H. Sit, M. S. Jalal, and R. Asmus. The authors thank A. Earle and J. Lammerding for assistance in maintaining the mdx mouse colony. The authors thank R. Puri from the Cornell Progressive Assessment of Therapeutics (PATh) Facility for maintaining the NSG mouse colony.

## Author contributions

A.M.L., K.H.K., S.Y.S., V.M.A., and B.D.C. designed the study. A.M.L., S.Y.S., V.M.A., P.F., and E.H.H.F. organized the mouse colony and performed animal procedures and cell isolations. V.M.A. and B.D.C., with assistance from R.M. and L.A.A., performed hydrogel characterization studies. A.M.L., K.H.K., S.Y.S., and V.M.A. conducted the long-term cell culture studies. K.H.K., V.M.A., and P.F. performed and analyzed the gene expression studies. A.M.L. and R.F.K. performed and analyzed the immunoblot assays. K.H.K., A.M.L., and J.C.C. conducted the Pax7-zsGreen imaging studies. K.H.K., S.Y.S., P.F., E.H.H.F., and B.D.C. performed the cell transplantation studies. A.M.L., K.H.K., and V.M.A. performed the statistical analyses. A.M.L., K.H.K., V.M.A., and B.D.C. wrote the manuscript. All authors reviewed the manuscript.

## Author contributions using CRediT taxonomy

Conceptualization and Methodology, A.M.L., K.H.K., S.Y.S., V.M.A., B.D.C.;

Investigation and Formal Analysis, A.M.L., K.H.K., S.Y.S., V.M.A., J.C.C., R.F.K., P.F., E.H.H.F, R.M.;

Writing – Original Draft, A.M.L., K.H.K., V.M.A., B.D.C.; Writing – Review and Editing, A.M.L., B.D.C.;

Funding Acquisition and Supervision, B.D.C., L.A.A.

## Supplementary Figures and Legends

**Figure S1.**
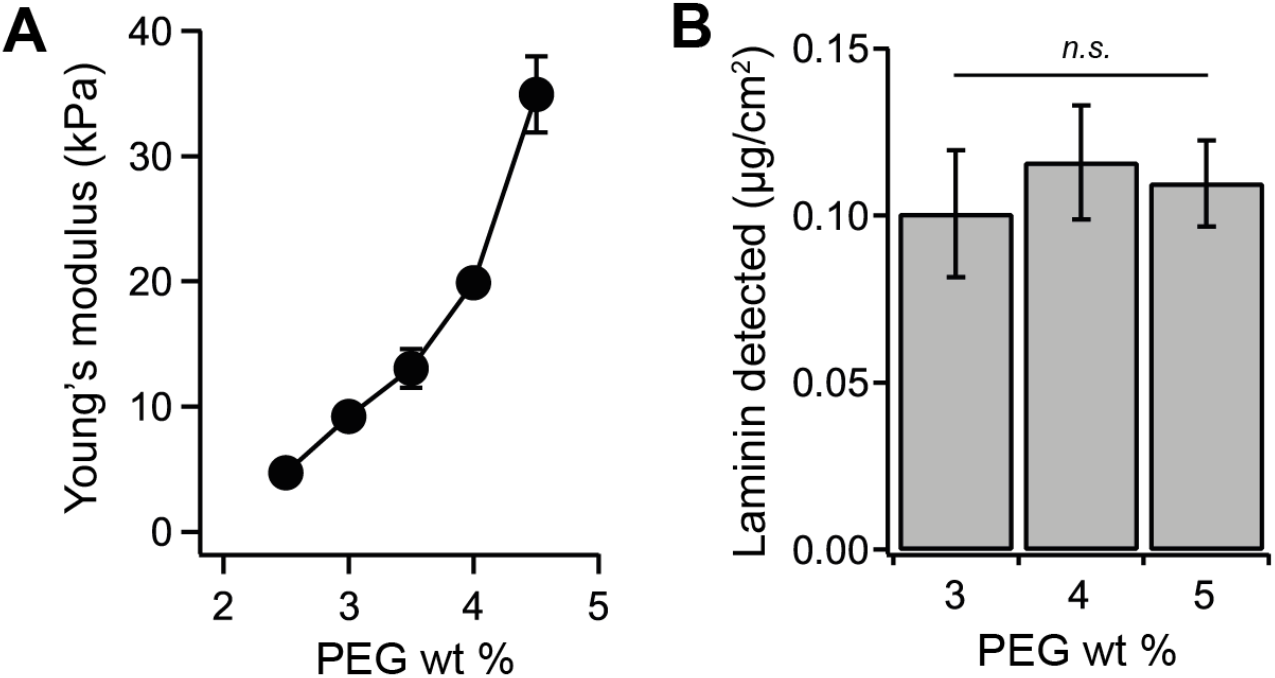
Biophysical and biomolecular characterization of laminin-conjugated PEG hydrogels with varying PEG weight percentage (Related to Figure 1). (**A**) Bulk Young’s modulus of PEG hydrogels, quantified by shear rheometry, synthesized with 2.5-4.5 weight percentage (wt%) total PEG. Mean ± s.e.m., n = 3. **(B**) Laminin concentration detected on PEG hydrogels via immunodetection and secondary chemiluminescence. Mean ± s.e.m., n = 3. * denotes *P* < 0.05 or not significant (n.s.) by Student’s T-test in (**B**).

**Figure S2.**
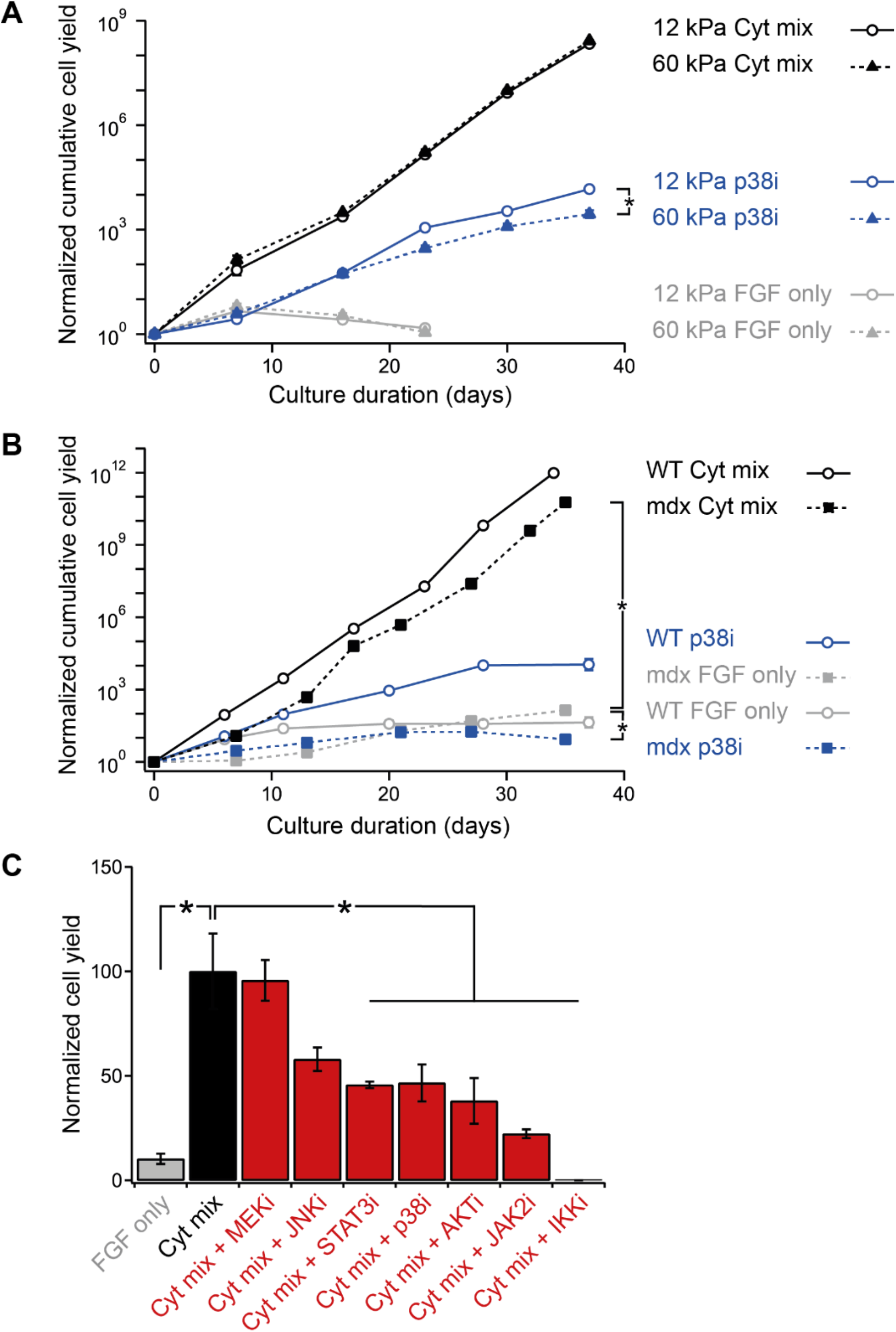
Short-term and long-term MuSC proliferation in response to cytokine mix treatment (Related to Figure 2). (**A-B**) Long-term MuSC cultures on hydrogels passaged every 6-8 d and treated with FGF or FGF + cytokine mix (TNF-α, IL-1α, IL-13, and IFN-γ) or p38i (SB203580, 5 μM) for 5 wk. All cell counts normalized to each seeded cell at d 0. (**A**) Normalized cumulative cell yield counts from 0-40 d for C57BL/6J WT MuSCs treated with (i) FGF only, (ii) FGF + cytokine mix, and (iii) FGF + p38i on 12 kPa or 60 kPa hydrogels. Mean ± s.e.m., n = 4. (**B**) Normalized cumulative cell yield counts from 0-40 d for C57BL/6J WT vs Dmd^mdx^ (mdx) MuSCs treated with (i) FGF only, (ii) FGF + cytokine mix, and (iii) FGF + p38i on 12 kPa hydrogels. Mean ± s.e.m., n = 4. (**C**) Short-term cultures of WT MuSC on 12 kPa hydrogels and treated with FGF2 or cytokine mix, with or without a small molecule inhibitor (MEKi: PD0325901, 10 nM; JNKi: SP600125, 1 μM; STAT3i: 5,15-DPP, 5 μM; p38i: SB203580, 5 μM; AKTi: GSK2110183, 10 nM; JAK2i: AG-490, 1 μM; IKKi: BMS-345541, 10 μM). Normalized cell yield at 1 wk. Mean ± s.e.m., n = 3. * denotes *P* < 0.05 by Student’s T-test for log-transformed values at the 5-wk timepoint in (**A-B**) and for non-transformed values in (**C**).

**Figure S3.**
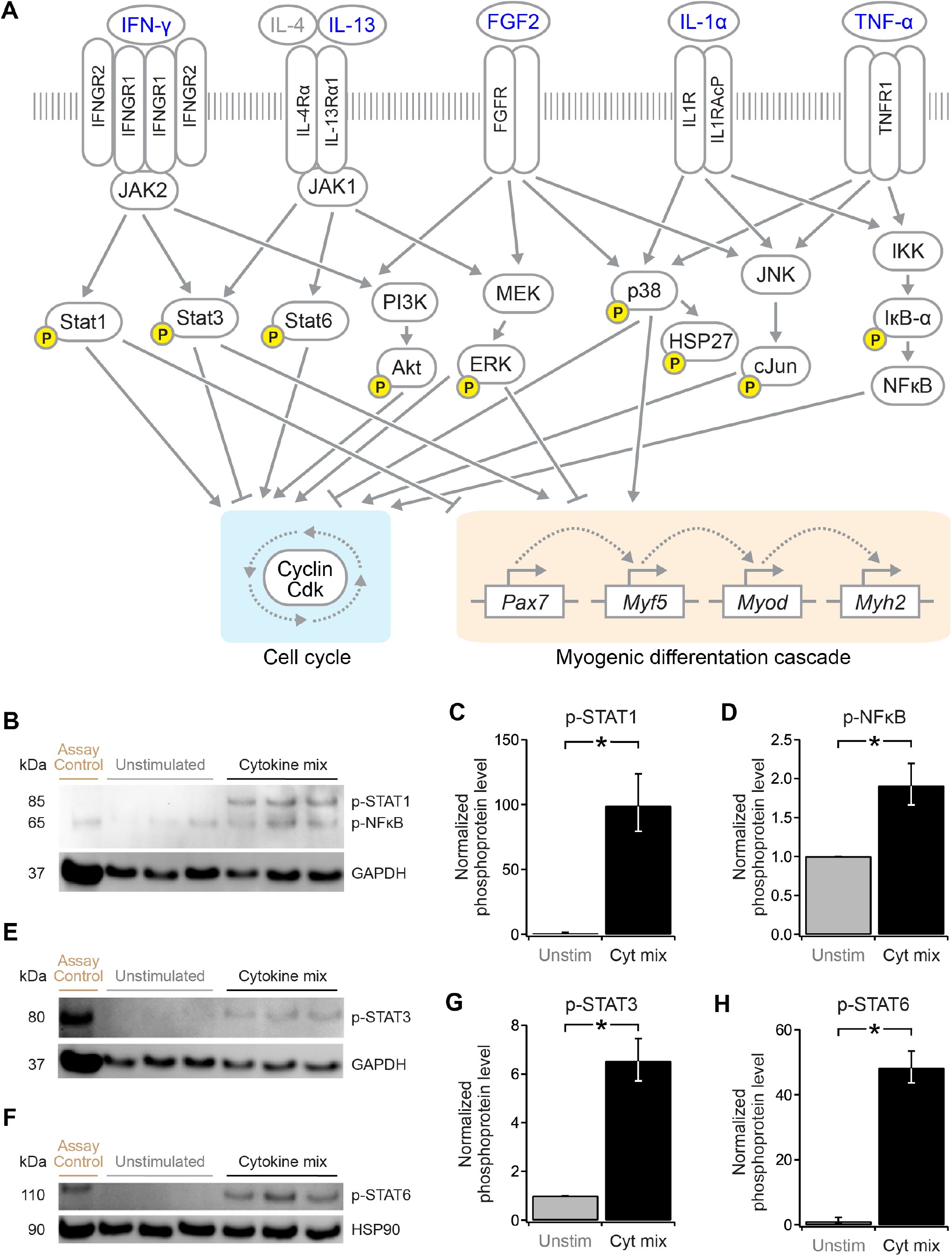
Cytokine treatment induces intracellular signaling activation (Related to Figure 3). (**A**) Pathway diagram displaying known activating and inhibiting effects of FGF2 and the TNF-α, IL-1α, IL-13, and IFN-γ cytokine mix factors on numerous intracellular signaling cascades, cell cycle genes, and myogenic regulatory genes. Yellow “p” indicates phosphoprotein mediator measured by Luminex in **Fig. 3**. (**B-H**) Primary myoblasts were cultured on 12 kPa hydrogels in growth medium and then switched to basal medium and were stimulated with the cytokine mix for 30 min (or left unstimulated) and then were lysed and analyzed by immunoblotting. (**B, E, F**) Immunoblots for STAT1 (pY701), NFκB (pS536), STAT3 (pY705), STAT6 (pY641), GAPDH, and/or HSP90. (**C, D, G, H**) Phosphoprotein band abundance normalized to GAPDH or HSP90 as a housekeeping protein, and then averaged across biological replicates from (**B, E, F**). Mean ± s.e.m., n = 3. * denotes *P* < 0.05 by Student’s T-test.

**Figure S4.**
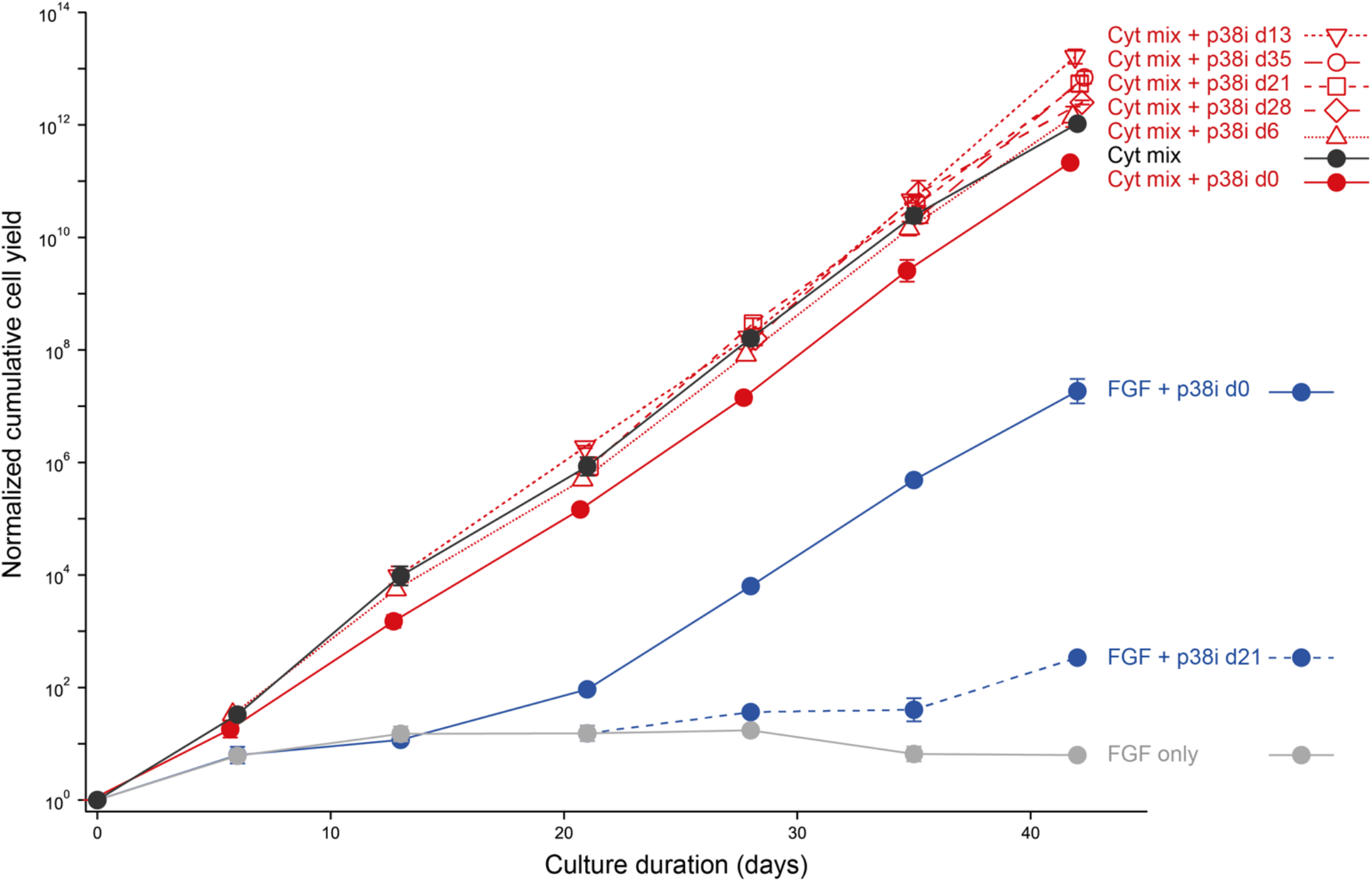
Late-stage inhibition of p38α/β MAPK enhances MuSC expansion in long-term cultures (Related to Figure 4). Long-term MuSC cultures on 12 kPa laminin-conjugated hydrogels were passaged every 6-8 d and treated with FGF or FGF + cytokine mix (TNF-α, IL-1α, IL-13, IFN-γ), without or with p38i (SB203580, 5 μM; addition staged weekly). “p38i d*X*” indicates SB203580 was added starting at day *X* and maintained in every media change thereafter. Normalized cumulative cell yields from 0-42 d. Mean ± s.e.m., n = 4. Statistical analyses reported for 42 d timepoint in **Fig. 4B**.

**Figure S5.**
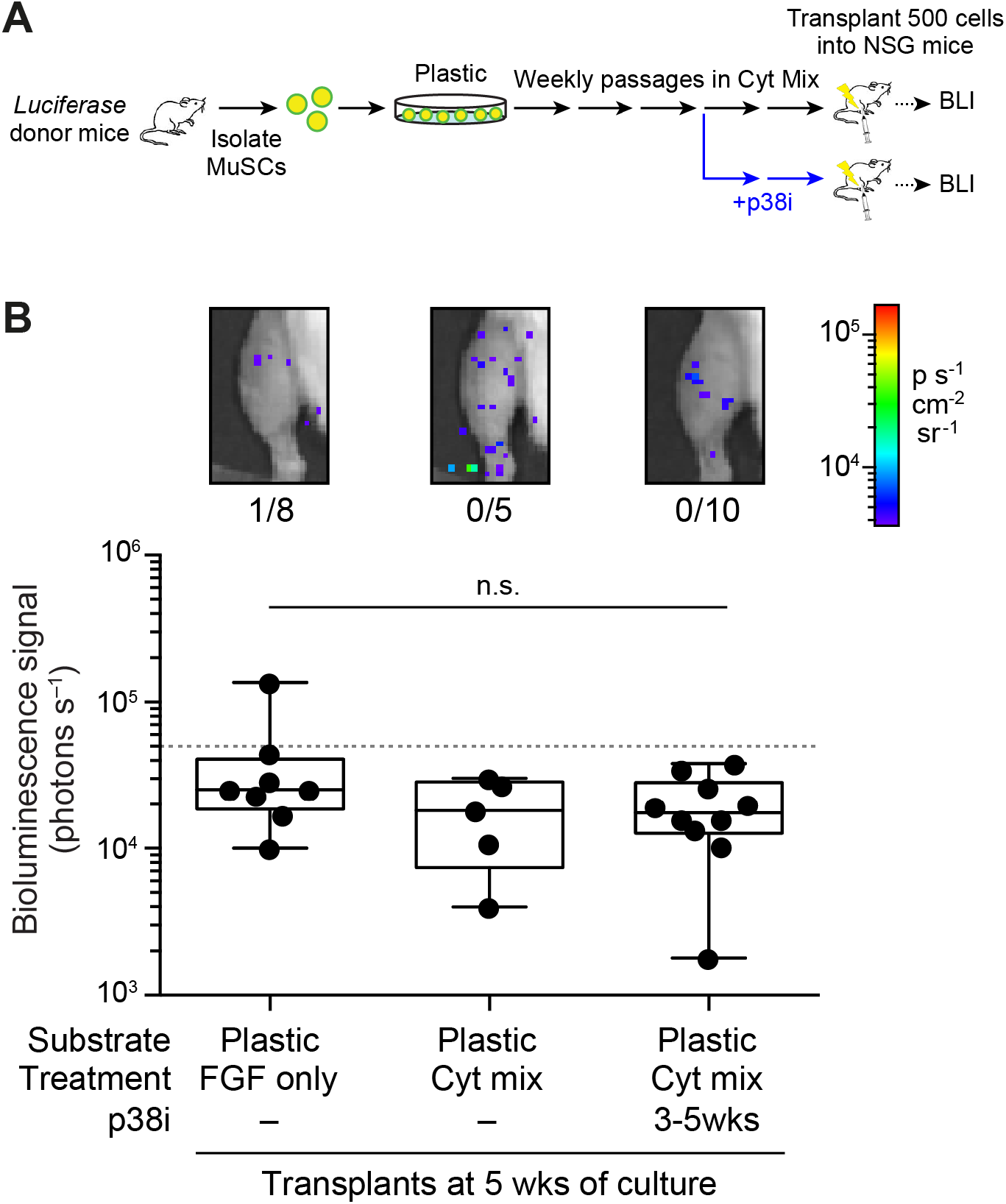
Plastic substrates are not permissive to enhancement of transplant engraftment potential by late-phase p38i treatment in long-term culture (Related to Figure 6). (**A**) Transplant assay schematic. Luciferase-expressing MuSCs were cultured and passaged weekly on laminin-conjugated tissue culture plastic substrates in the presence of FGF +/- cytokine mix mix (TNF-α, IL-1α, IL-13, IFN-γ), without or with p38i starting at 3 wk. At 5 wk of culture, 500 cells were transplanted into NSG mice and engraftment was measured using bioluminescent imaging (BLI) at one-month post-transplantation. (**B**) Background-subtracted representative BLI images (top) and signals (bottom; n = 5-10) at one-month post-transplantation for each culture condition are reported. Dashed line indicates positive engraftment threshold of 50,000 photons s^−1^. Fraction of transplants resulting in positive engraftment outcome reported in inset. All comparisons were not significant (n.s.) by Mann-Whitney U test.

## References

Almada, A.E., and Wagers, A.J. (2016). Molecular circuitry of stem cell fate in skeletal muscle regeneration, ageing and disease. Nat. Rev. Mol. Cell Biol. 17, 267–279.

Arpke, R.W., Darabi, R., Mader, T.L., Zhang, Y., Toyama, A., Lonetree, C.L., Nash, N., Lowe, D.A., Perlingeiro, R.C.R., and Kyba, M. (2013). A new immuno-, dystrophin-deficient model, the NSG-mdx(4Cv) mouse, provides evidence for functional improvement following allogeneic satellite cell transplantation. Stem Cells 31, 1611–1620.

Bentzinger, C.F., Wang, Y.X., Dumont, N.A., and Rudnicki, M.A. (2013). Cellular dynamics in the muscle satellite cell niche. EMBO Rep. 14, 1062–1072.

Bernet, J.D., Doles, J.D., Hall, J.K., Kelly Tanaka, K., Carter, T.A., and Olwin, B.B. (2014). p38 MAPK signaling underlies a cell-autonomous loss of stem cell self-renewal in skeletal muscle of aged mice. Nat. Med. 20, 265–271.

Blau, H.M., and Webster, C. (1981). Isolation and characterization of human muscle cells. Proc. Natl. Acad. Sci. U. S. A. 78, 5623–5627.

Blau, H.M., Cosgrove, B.D., and Ho, A.T.V. (2015). The central role of muscle stem cells in regenerative failure with aging. Nat. Med. 21, 854–862.

Borselli, C., Cezar, C.A., Shvartsman, D., Vandenburgh, H.H., and Mooney, D.J. (2011). The role of multifunctional delivery scaffold in the ability of cultured myoblasts to promote muscle regeneration. Biomaterials 32, 8905–8914.

Bosnakovski, D., Xu, Z., Li, W., Thet, S., Cleaver, O., Perlingeiro, R.C., and Kyba, M. (2008). Prospective isolation of skeletal muscle stem cells with a Pax7 reporter. Stem Cells 26, 3194–3204.

Bouchentouf, M., Skuk, D., and Tremblay, J.P. (2007). Early and massive death of myoblasts transplanted into skeletal muscle: Responsible factors and potential solutions. Curr. Opin. Organ Transplant. 12, 664–667.

Brien, P., Pugazhendhi, D., Woodhouse, S., Oxley, D., and Pell, J.M. (2013). p38alpha MAPK regulates adult muscle stem cell fate by restricting progenitor proliferation during postnatal growth and repair. Stem Cells 31, 1597–1610.

Carraher Jr., C.E. (2016). Carraher’s Polymer Chemistry.

Castro, F., Cardoso, A.P., Gonçalves, R.M., Serre, K., and Oliveira, M.J. (2018). Interferon-Gamma at the Crossroads of Tumor Immune Surveillance or Evasion. Front. Immunol. 9, 847.

Charville, G.W., Cheung, T.H., Yoo, B., Santos, P.J., Lee, G.K., Shrager, J.B., and Rando, T.A. (2015). Ex Vivo Expansion and In Vivo Self-Renewal of Human Muscle Stem Cells. Stem Cell Reports 5, 621–632.

Chen, S.E., Jin, B., and Li, Y.P. (2007). TNF-α regulates myogenesis and muscle regeneration by activating p38 MAPK. Am. J. Physiol. Cell Physiol. 292, C1660–C1671.

Collinsworth, A.M., Zhang, S., Kraus, W.E., and Truskey, G.A. (2002). Apparent elastic modulus and hysteresis of skeletal muscle cells throughout differentiation. Am. J. Physiol. Cell Physiol. 283, C1219–27.

Cosgrove, B.D., Sacco, A., Gilbert, P.M., and Blau, H.M. (2009). A home away from home: challenges and opportunities in engineering in vitro muscle satellite cell niches. Differentiation 78, 185–194.

Cosgrove, B.D., Gilbert, P.M., Porpiglia, E., Mourkioti, F., Lee, S.P., Corbel, S.Y., Llewellyn, M.E., Delp, S.L., and Blau, H.M. (2014). Rejuvenation of the muscle stem cell population restores strength to injured aged muscles. Nat. Med. 20, 255–264.

Davoudi, S., Chin, C.Y., Cooke, M.J., Tam, R.Y., Shoichet, M.S., and Gilbert, P.M. (2018). Muscle stem cell intramuscular delivery within hyaluronan methylcellulose improves engraftment efficiency and dispersion. Biomaterials 173, 34–46.

Egerman, M.A., and Glass, D.J. (2014). Signaling pathways controlling skeletal muscle mass. Crit. Rev. Biochem. Mol. Biol. 49, 59–68.

Engler, A.J., Griffin, M.A., Sen, S., Bonnemann, C.G., Sweeney, H.L., and Discher, D.E. (2004). Myotubes differentiate optimally on substrates with tissue-like stiffness: pathological implications for soft or stiff microenvironments. J. Cell Biol. 166, 877–887.

Engler, A.J., Sen, S., Sweeney, H.L., and Discher, D.E. (2006). Matrix elasticity directs stem cell lineage specification. Cell 126, 677–689.

Fu, X., Xiao, J., Wei, Y., Li, S., Liu, Y., Yin, J., Sun, K., Sun, H., Wang, H., Zhang, Z., et al. (2015). Combination of inflammation-related cytokines promotes long-term muscle stem cell expansion. Cell Res. 25, 1082–1083.

Gao, Y., Kostrominova, T.Y., Faulkner, J.A., and Wineman, A.S. (2008). Age-related changes in the mechanical properties of the epimysium in skeletal muscles of rats. J. Biomech. 41, 465–469.

Gilbert, P.M., Havenstrite, K.L., Magnusson, K.E.G., Sacco, A., Leonardi, N.A., Kraft, P., Nguyen, N.K., Thrun, S., Lutolf, M.P., and Blau, H.M. (2010). Substrate Elasticity Regulates Skeletal Muscle Stem Cell Self-Renewal in Culture. Science (80-.). 329, 1078–1081.

Le Grand, F., Jones, A.E., Seale, V., Scime, A., Rudnicki, M.A., Scimè, A., and Rudnicki, M.A. (2009). Wnt7a activates the planar cell polarity pathway to drive the symmetric expansion of satellite stem cells. Cell Stem Cell 4, 535–547.

Gussoni, E., Pavlath, G.K., Lanctot, A.M., Sharma, K.R., Miller, R.G., Steinman, L., and Blau, H.M. (1992). Normal dystrophin transcripts detected in Duchenne muscular dystrophy patients after myoblast transplantation. Nature 356, 435–438.

Gussoni, E., Blau, H.M., and Kunkel, L.M. (1997). The fate of individual myoblasts after transplantation into muscles of DMD patients. Nat. Med. 3, 970–977.

Han, W.M., Anderson, S.E., Mohiuddin, M., Barros, D., Nakhai, S.A., Shin, E., Amaral, I.F., Pêgo, A.P., García, A.J., and Jang, Y.C. (2018). Synthetic matrix enhances transplanted satellite cell engraftment in dystrophic and aged skeletal muscle with comorbid trauma. Sci. Adv. 4.

Heredia, J.E., Mukundan, L., Chen, F.M., Mueller, A.A., Deo, R.C., Locksley, R.M., Rando, T.A., and Chawla, A. (2013). Type 2 innate signals stimulate fibro/adipogenic progenitors to facilitate muscle regeneration. Cell 153, 376–388.

Jones, N.C., Tyner, K.J., Nibarger, L., Stanley, H.M., Cornelison, D.D.W., Fedorov, Y. V., and Olwin, B. (2005). The p38α/β MAPK functions as a molecular switch to activate the quiescent satellite cell. J. Cell Biol. 169, 105–116.

Judson, R.N., and Rossi, F.M.V. (2020). Towards stem cell therapies for skeletal muscle repair (Nature Research).

Judson, R.N., Quarta, M., Oudhoff, M.J., Soliman, H., Yi, L., Chang, C.K., Loi, G., Vander Werff, R., Cait, A., Hamer, M., et al. (2018). Inhibition of Methyltransferase Setd7 Allows the In Vitro Expansion of Myogenic Stem Cells with Improved Therapeutic Potential. Cell Stem Cell 22, 177–190.e7.

Lean, G., Halloran, M.W., Marescal, O., Jamet, S., Lumb, J.P., and Crist, C. (2019). Ex vivo Expansion of Skeletal Muscle Stem Cells with a Novel Small Compound Inhibitor of eIF2α Dephosphorylation. Regen. Med. Front. 1, e190003.

Li, W., Moylan, J.S., Chambers, M.A., Smith, J., and Reid, M.B. (2009). Interleukin-1 stimulates catabolism in C2C12 myotubes. Am. J. Physiol. Cell Physiol. 297, C706–C714.

Lluis, F., Perdiguero, E., Nebreda, A.R., and Munoz-Canoves, P. (2006). Regulation of skeletal muscle gene expression by p38 MAP kinases. Trends Cell Biol. 16, 36–44.

Loiben, A.M., Soueid-Baumgarten, S., Kopyto, R.F., Bhattacharya, D., Kim, J.C., and Cosgrove, B.D. (2017). Data-Modeling Identifies Conflicting Signaling Axes Governing Myoblast Proliferation and Differentiation Responses to Diverse Ligand Stimuli. Cell. Mol. Bioeng. 10, 433–450.

Lutolf, M.P., and Hubbell, J.A. (2003). Synthesis and physicochemical characterization of end-linked poly(ethylene glycol)-co-peptide hydrogels formed by Michael-type addition. Biomacromolecules 4, 713–722.

Lutolf, M.P., Gilbert, P.M., and Blau, H.M. (2009). Designing materials to direct stem-cell fate. Nature 462, 433–441.

McCormick, S.M., and Heller, N.M. (2015). Commentary: IL-4 and IL-13 receptors and signaling. Cytokine 75, 38–50.

De Micheli, A.J., Laurilliard, E.J., Heinke, C.L., Ravichandran, H., Fraczek, P., Soueid-Baumgarten, S., De Vlaminck, I., Elemento, O., and Cosgrove, B.D. (2020). Single-Cell Analysis of the Muscle Stem Cell Hierarchy Identifies Heterotypic Communication Signals Involved in Skeletal Muscle Regeneration. Cell Rep. 30, 3583–3595.e5.

Montarras, D., Morgan, J., Colins, C., Relaix, F., Zaffran, S., Cumano, A., Partridge, T., Buckingham, M., Collins, C., Relaix, F., et al. (2005). Direct isolation of satellite cells for skeletal muscle regeneration. Science (80-.). 309, 2064–2067.

Morrissey, J.B., Cheng, R.Y., Davoudi, S., and Gilbert, P.M. (2016). Biomechanical Origins of Muscle Stem Cell Signal Transduction. J. Mol. Biol 428, 1441–1454.

Palacios, D., Mozzetta, C., Consalvi, S., Caretti, G., Saccone, V., Proserpio, V., Marquez, V.E., Valente, S., Mai, A., Forcales, S. V, et al. (2010). TNF/p38alpha/polycomb signaling to Pax7 locus in satellite cells links inflammation to the epigenetic control of muscle regeneration. Cell Stem Cell 7, 455–469.

Pawlikowski, B., Vogler, T.O., Gadek, K., and Olwin, B.B. (2017). Regulation of skeletal muscle stem cells by fibroblast growth factors. Dev. Dyn. 246, 359–367.

Perdiguero, E., Ruiz-Bonilla, V., Gresh, L., Hui, L., Ballestar, E., Sousa-Victor, P., Baeza-Raja, B., Jardi, M., Bosch-Comas, A., Esteller, M., et al. (2007). Genetic analysis of p38 MAP kinases in myogenesis: fundamental role of p38alpha in abrogating myoblast proliferation. EMBO J. 26, 1245–1256.

Price, F.D., Kuroda, K., and Rudnicki, M.A. (2007). Stem cell based therapies to treat muscular dystrophy. Biochim. Biophys. Acta 1772, 272–283.

Price, F.D., von Maltzahn, J., Bentzinger, C.F., Dumont, N.A., Yin, H., Chang, N.C., Wilson, D.H., Frenette, J., and Rudnicki, M.A. (2014). Inhibition of JAK-STAT signaling stimulates adult satellite cell function. Nat. Med. 20, 1174–1181.

Qing, Y., and Stark, G.R. (2004). Alternative activation of STAT1 and STAT3 in response to interferon- γ. J. Biol. Chem. 279, 41679–41685.

Quarta, M., Brett, J.O., DiMarco, R., De Morree, A., Boutet, S.C., Chacon, R., Gibbons, M.C., Garcia, V.A., Su, J., Shrager, J.B., et al. (2016). An artificial niche preserves the quiescence of muscle stem cells and enhances their therapeutic efficacy. Nat. Biotechnol. 34, 752–759.

Rando, T.A., and Blau, H.M. (1994). Primary mouse myoblast purification, characterization, and transplantation for cell-mediated gene therapy. J. Cell Biol. 125, 1275–1287.

Rao, N., Agmon, G., Tierney, M.T., Ungerleider, J.L., Braden, R.L., Sacco, A., and Christman, K.L. (2017). Engineering an Injectable Muscle-Specific Microenvironment for Improved Cell Delivery Using a Nanofibrous Extracellular Matrix Hydrogel. ACS Nano 11, 3851–3859.

Rinaldi, F., and Perlingeiro, R.C. (2014). Stem cells for skeletal muscle regeneration: therapeutic potential and roadblocks. Transl. Res. 163, 409–417.

Rosant, C., Nagel, M.D., and Perot, C. (2007). Aging affects passive stiffness and spindle function of the rat soleus muscle. Exp. Gerontol. 42, 301–308.

Sacco, A., Doyonnas, R., Kraft, P., Vitorovic, S., and Blau, H.M. (2008). Self-renewal and expansion of single transplanted muscle stem cells. Nature 456, 502–506.

Sampath, S.C., Sampath, S.C., Ho, A.T. V., Corbel, S.Y., Millstone, J.D., Lamb, J., Walker, J., Kinzel, B., Schmedt, C., and Blau, H.M. (2018). Induction of muscle stem cell quiescence by the secreted niche factor Oncostatin M. Nat. Commun. 9, 1531.

Sanes, J.R. (2003). The basement membrane/basal lamina of skeletal muscle. J. Biol. Chem. 278, 12601–12604.

Schroder, K., Hertzog, P.J., Ravasi, T., and Hume, D.A. (2004). Interferon-γ: an overview of signals, mechanisms and functions. J. Leukoc. Biol. 75, 163–189.

Segalés, J., Islam, A.B.M.M.K., Kumar, R., Liu, Q.-C., Sousa-Victor, P., Dilworth, F.J., Ballestar, E., Perdiguero, E., and Muñoz-Cánoves, P. (2016). Chromatin-wide and transcriptome profiling integration uncovers p38α MAPK as a global regulator of skeletal muscle differentiation. Skelet. Muscle 6, 9.

Skuk, D. (2004). Myoblast transplantion for inherited myopathies: A clinical approach. Expert Opin. Biol. Ther. 4, 1871–1885.

Sleep, E., Cosgrove, B.D., McClendon, M.T., Preslar, A.T., Chen, C.H., Sangji, M.H., Pérez, C.M.R., Haynes, R.D., Meade, T.J., Blau, H.M., et al. (2017). Injectable biomimetic liquid crystalline scaffolds enhance muscle stem cell transplantation. Proc. Natl. Acad. Sci. U. S. A. 114, E7919–E7928.

Stedman, H.H., Sweeney, H.L., Shrager, J.B., Maguire, H.C., Panettieri, R.A., Petrof, B., Narusawa, M., Leferovich, J.M., Sladky, J.T., and Kelly, A.M. (1991). The mdx mouse diaphragm reproduces the degenerative changes of Duchenne muscular dystrophy. Nature 352, 536–539.

Tidball, J.G. (2017). Regulation of muscle growth and regeneration by the immune system. Nat. Rev. Immunol. 17, 165–178.

Tierney, M.T., Aydogdu, T., Sala, D., Malecova, B., Gatto, S., Puri, P.L., Latella, L., and Sacco, A. (2014). STAT3 signaling controls satellite cell expansion and skeletal muscle repair. Nat. Med. 20, 1182–1186.

Wang, Y.X., and Rudnicki, M.A. (2012). Satellite cells, the engines of muscle repair. Nat. Rev. Mol. Cell Biol. 13, 127–133.

Wolf, M.T., Dearth, C.L., Sonnenberg, S.B., Loboa, E.G., and Badylak, S.F. (2015). Naturally derived and synthetic scaffolds for skeletal muscle reconstruction. Adv. Drug Deliv. Rev. 84, 208–221.

Wosczyna, M.N., and Rando, T.A. (2018). A Muscle Stem Cell Support Group: Coordinated Cellular Responses in Muscle Regeneration. Dev Cell 46, 135–143.

Xu, C., Tabebordbar, M., Iovino, S., Ciarlo, C., Liu, J., Castiglioni, A., Price, E., Liu, M., Barton, E.R., Kahn, C.R., et al. (2013). A Zebrafish Embryo Culture System Defines Factors that Promote Vertebrate Myogenesis across Species. Cell 155, 909–921.

Yablonka-Reuveni, Z., Seger, R., and Rivera, A.J. (1999). Fibroblast growth factor promotes recruitment of skeletal muscle satellite cells in young and old rats. J. Histochem. Cytochem. 47, 23–42.

Yang, W., and Hu, P. (2018). Skeletal muscle regeneration is modulated by inflammation. J. Orthop. Transl. 13, 25–32.

Yin, H., Price, F., and Rudnicki, M.A. (2013). Satellite Cells and the Muscle Stem Cell Niche. Physiol. Rev. 93, 23–67.

Zismanov, V., Chichkov, V., Colangelo, V., Jamet, S., Wang, S., Syme, A., Koromilas, A.E., and Crist, C. (2016). Phosphorylation of eIF2α is a Translational Control Mechanism Regulating Muscle Stem Cell Quiescence and Self-Renewal. Cell Stem Cell 18, 79–90.

